# Non-invasive peripheral delivery of CDNF fragment protects neurons in models of Parkinson’s and ALS

**DOI:** 10.1101/2025.02.09.637291

**Authors:** Li Ying Yu, Tuulikki Viljakainen, Liam Beckett, Jinhan Nam, Anu Hyysalo, Patrick Lüningschrör, Katrina Albert, Tapani Koppinen, Tanja Hyvärinen, Helena Tossavainen, Maarit Hellman, Mikko Airavaara, Michael Sendtner, Perttu Permi, Päivi Lindholm, Mart Saarma, Merja H. Voutilainen

## Abstract

Non-invasive delivery of brain therapeutics is a key challenge for treating neurodegenerative diseases. Here, we discovered a novel carboxy (C)-terminal fragment of cerebral dopamine neurotrophic factor (C-CDNF) that protects dopamine (DA) and motoneurons (MNs) in rodent models of Parkinson’s disease (PD) and amyotrophic lateral sclerosis (ALS). C-CDNF retains the same structure as CDNF and similarly to CDNF regulates cell stress pathways but unorthodoxly enters cultured neurons and passes through the blood-brain barrier. *In vivo*, intracranially or peripherally delivered C-CDNF improves motor deficits, protects DA neurons, and restores motor behavior in a rat model of PD. Subcutaneous C-CDNF also protects MNs and reduces microglial activation in an ALS model. Based on our findings, beginning C-CDNF treatment soon after diagnosis is anticipated to delay progression of PD and ALS, thereby improving treatment outcome. Thus, systemic delivery of C-CDNF should simplify the administration of protein-based therapeutics to patients while reducing treatment risk and financial burden for patients and families.

## Introduction

Neurodegenerative disorders including Parkinsońs disease (PD) and amyotrophic lateral sclerosis (ALS) are incurable conditions and eventually leading to death. There is an unmet need for new and more effective treatments that could slow down or even stop disease progression. Neurodegenerative diseases are characterized by the vulnerability of a specific neural population, that gives rise to a specific set of symptoms. Nevertheless, the pathogeneses of neurodegenerative diseases, such as ALS and PD, show striking similarities. Common mechanisms observed in many neurodegenerative diseases include inflammation, mitochondrial dysfunction, reduced growth factor levels, apoptosis, and endoplasmic reticulum (ER) stress. Accumulation of misfolded and aggregated proteins in the ER disrupt proteostasis, subsequently triggering ER stress.

The unfolded protein response (UPR), a cellular defence signalling cascade, alleviates ER stress by; (i) suppressing mRNA translation in ribosomes, (ii) degrading misfolded proteins, and (iii) activating signalling pathways that lead to the production of molecular chaperones that facilitate protein folding. UPR activation in mammalian cells is mediated by three ER transmembrane receptors, including IRE1α (inositol-requiring enzyme 1α), PERK (PKR-like ER kinase), and ATF6 (activating transcription factor 6). Acute UPR activation is initially protective. However, under conditions of prolonged ER stress, the UPR promotes apoptosis, eventually leading to cell death (Szegezdi et al. 2006; Voutilainen et al. 2015; De Lorenzo et al. 2023). Chronic ER stress is associated with many pathophysiological conditions, including PD and ALS (Hetz and Mollereau 2014).

Neurotrophic factors (NTFs) such as glial cell line-derived neurotrophic factor (GDNF) and brain-derived neurotrophic factor (BDNF) are secretory proteins that regulate the development and maintenance of neurons, neurite growth, and neuroregeneration (Airaksinen and Saarma 2002). They have been explored as novel drug candidates for the treatment of PD and ALS but their efficacy in clinical trials has been modest (Thoenen and Sendtner 2002; Barker et al. 2020), mostly due to adverse pharmacokinetic properties. Cerebral dopamine neurotrophic factor (CDNF), and mesencephalic astrocyte-derived neurotrophic factor (MANF) are proteins with NTF-like properties (Lindholm et al. 2007; Petrova et al. 2003; Voutilainen et al. 2009; Voutilainen et al. 2011; Lindholm and Saarma 2022).

CDNF is a small protein (18kDa, 161 aa) that contains a signal sequence for secretion and an ER retention signal at the C-terminus (Lindholm et al. 2007; Liu et al. 2015; Norisada et al. 2016; Galli et al. 2019)**(Fig 1A-C)**. CDNF is mainly located in the ER (Eesmaa et al. 2022) but is also secreted under conditions of cellular stress and is found in human serum (Galli et al. 2019). Structural studies show that it consists of two independent domains: a saposin-like N-terminal domain and a C-terminal SAP-like domain (Hellman et al. 2011; Latge et al. 2015; Parkash et al. 2009)**(Fig. 1A-C)**. The C-terminal domain is potently anti-apoptotic (Hellman et al. 2011; Lindholm and Saarma 2022). CDNF differs from known NTFs in having a unique dual mode of action (Parkash et al. 2009; Eesmaa et al. 2022). During cellular homeostasis, it is located primarily in the luminal side of the ER and regulates UPR and ER stress intracellularly (Matlik et al. 2015; Eesmaa et al. 2022). CDNF interacts with the ER chaperone GRP78 (also known as BiP) (Eesmaa et al. 2022). CDNF is upregulated and secreted in response to experimental ER stress and protects cells from ER stress-induced cell death (De Lorenzo et al. 2023; Eesmaa et al. 2022). Extracellularly, and *in vivo*, CDNF acts only on ER stressed and injured cells and inhibits the synthesis and release of pro-inflammatory cytokines (Nadella et al. 2014; Tseng et al. 2023; Zhao et al. 2016). CDNF cellular effects can be blocked by IRE1α and PERK inhibitors strongly suggesting the involvement of UPR receptors in CDNF signalling (De Lorenzo et al. 2023; Eesmaa et al. 2022).

**Figure 1.**
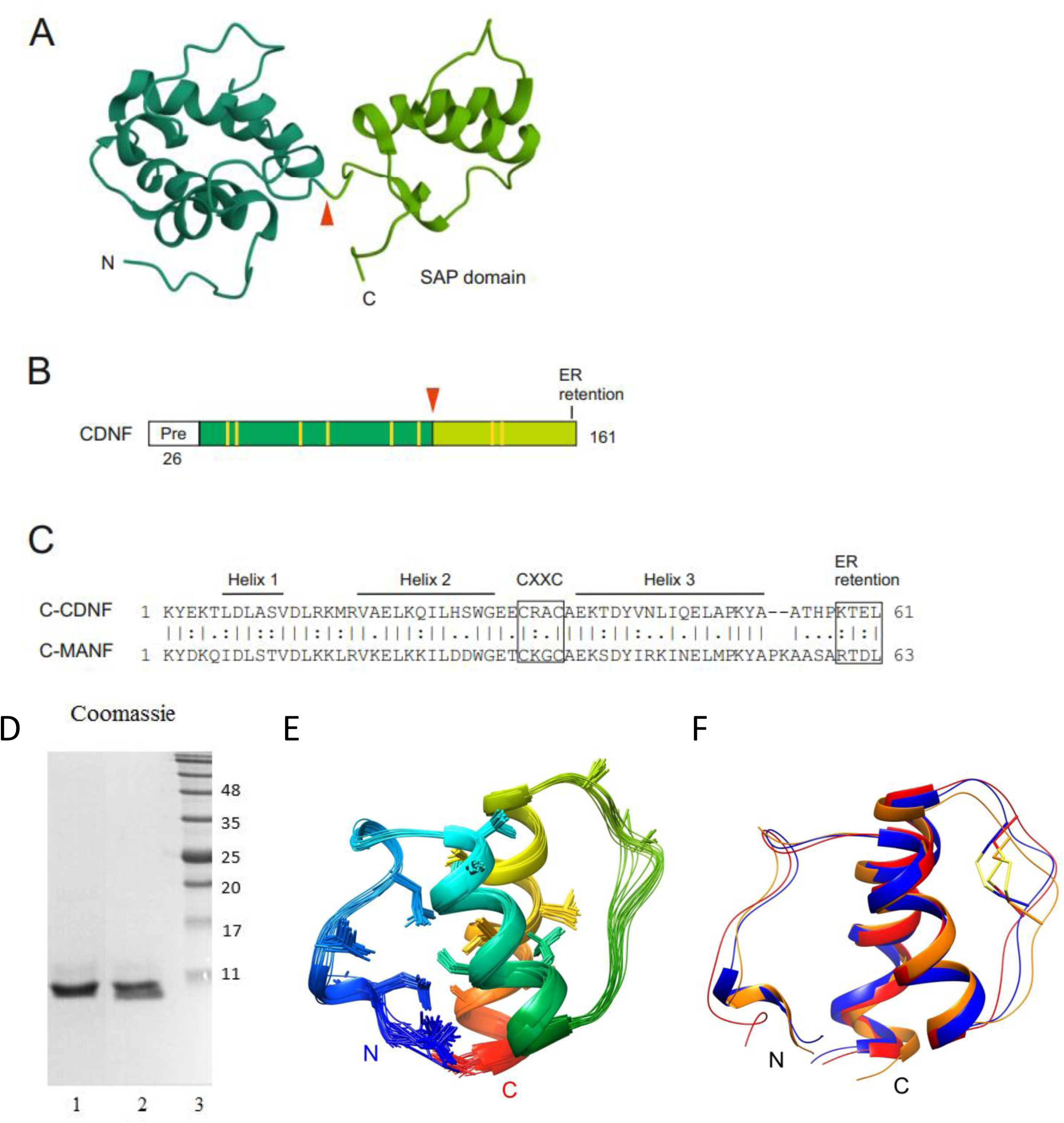
Structure and properties of C-CDNF. **A)** Structure of human CDNF PDB ID 4BIT, (Latge et al. 2015). The C-terminal SAP domain is indicated in lighter green. Arrowhead indicates the hypothetical cleavage site, releasing C-CDNF. The image was created from using Mol* 3D Viewer (Sehnal et al. 2021) on RCSB PDB (RCSB.org). **B)** Schematic primary structure of C-CDNF. Eight conserved cysteine residues are indicated in yellow. **C)** Alignment of C-CDNF and C-MANF amino acid sequences. Alpha-helical regions, CXXC motif and ER retention signal are indicated. **D**) SDS- polyacrylamide gel electrophoresis of CHO cell produced C-CDNF (2µg, lane 1), chemically synthesized C-CDNF (2 µg, lane 2), molecular weight markers (lane 3). **E**) Solution NMR structure of C-CDNF rainbow-colored from N- (blue) to C-terminus (red). To highlight structure precision, hydrophobic residues are shown in stick representation. Hydrogens as well as disordered N- and C- termini (residues 97-105, 154-161) have been omitted for clarity. **F**) Superimposition of C-CDNF (blue) with the same domain in CDNF (PDB ID 4BIT, orange) and C-MANF (2KVE, red). The disulphide bond is highlighted in the structure.

CDNF protects and restores the function of dopamine (DA) neurons in rodent and non-human primate (NHP) neurotoxin models of PD more effectively than other NTFs. Thus, it represents a compelling drug candidate for the disease-modifying treatment of PD (Lindholm et al. 2007; Voutilainen et al. 2011; Lindholm and Saarma 2022). Additionally, CDNF interacts directly with α-synuclein (α-syn), reducing its aggregation, neuronal entry, toxicity, and improves motor function in α-syn-based rodent models of PD (Albert et al. 2021).

CDNF has many beneficial properties compared to previously described NTFs: it does not affect naïve neurons, diffuses in brain tissue better than GDNF or BDNF, regulates the UPR and protein folding, and reduces inflammation (Voutilainen et al. 2011; De Lorenzo et al. 2023; Nadella et al. 2014; Tseng et al. 2023; Zhao et al. 2014). CDNF also increases motor coordination and protects motoneurons (MNs) by regulating ER stress in three different genetic rodent models of ALS (De Lorenzo et al. 2023). Based on the recent data from a phase I trial on PD patients, the primary endpoint was met: CDNF treatment was safe, and no side effects were observed (Huttunen et al. 2023). CDNF showed a therapeutic effect in some PD patients. However, in these clinical trials CDNF was injected into the brain using challenging stereotactic brain surgery. Since CDNF, GDNF, and other NTFs fail to cross the blood-brain barrier (BBB), they must be administered directly into the brain. In ALS clinical trials, NTFs have been delivered intrathecally or intracerebrally to patients (Barker et al., 2020; Thoenen and Sendtner 2002). In the recent clinical trials on PD patients CDNF and GDNF proteins were delivered to the caudate putamen (Huttunen et al. 2023; Whone et al. 2019; Barker et al. 2020). While promising, these invasive approaches bear significant risks and financial burden. For ethical reasons CDNF was tested in clinical trials in patients who had been diagnosed with moderate idiopathic PD in average for 10.5 years, a disease stage when 60-80% of the cell bodies and 70-90% of the axons of DA neurons have already been lost, thereby limiting the favourable outcome of neural protection while providing critical patient safety data.

Here, we discovered a novel brain-penetrating C-terminal fragment of CDNF (C-CDNF) that protects spinal moto- and nigral DA neurons *in vitro* and in animal models of ALS and PD. Brain-penetrating C-CDNF can be non-invasively delivered, thus C-CDNF is easy to use, safe, economical, more effective than invasive NTF treatments, and will enable the initiation of disease-modifying treatment immediately following the diagnosis of ALS and PD. Indeed, initiating treatment early in the course of the disease is anticipated to result in more effective treatment outcomes, since more neurons and neuronal circuits can be protected from degeneration. We envision that C-CDNF could be used as a basis for developing even more efficient brain penetrating therapeutic biomolecules, which alleviate ER stress and inflammation, reduce protein aggregation, protect, and regenerate neurons.

## Results

### Discovery of the CDNF C-terminal domain

We serendipitously discovered a new biologically active fragment of CDNF that we call C-CDNF. During routine purification of full-length recombinant human (rh)CDNF produced in insect Sf9 cells as described (Lindholm et al. 2007), we observed with immunoblotting that the purified CDNF preparation contained a smaller fragment (Saarma, Airavaara, Voutilainen et al. 2018). We initially assumed the fragment to be a proteolytic fragment of CDNF. However, the fragment was N-terminally sequenced and mass spectrometry (MS) revealed that it consists of amino acid residues 1- 107 of mature CDNF, i.e. being an N-terminal truncation of CDNF (Parkash et al. 2009) **(Fig. 1A)**. Thus, we reasoned that CDNF can be proteolytically cleaved into distinct N-terminal and C-terminal fragments. We expressed the N-terminal domain polypeptide of 100 amino acids (N-CDNF) and a 61 amino acid long C-terminal domain polypeptide (C-CDNF) in CHO cells **(Fig. 1A, B, D).** C-CDNF contains three alpha helices, a conserved CRAC sequence and the ER retention signal KTEL at the C-terminus **(Fig. 1C).** Each domain was expressed in CHO cells, purified and their amino acid sequences were verified by MS. Purified proteins were tested for their biological activity in several neuronal survival assays. We also studied the three-dimensional structure of C-CDNF using solution state NMR.

### C-terminal domain of CDNF retains the same structure as CDNF and is highly similar to C- MANF

To determine high-resolution solution NMR structure of C-CDNF, we produced uniformly ^15^N, ^13^C labelled polypeptide, corresponding to residues 101-161 in mature CDNF, using heterologous expression in *E. coli*. The isolated C-terminal domain adopts a structure **(Fig. 1E-F, Table S1)** very similar to that of the corresponding domain in the full-length CDNF (Latge et al. 2015); (PDB ID 4BIT, or the C-terminal domain of MANF (Hellman et al. 2011), (PDB ID 2KVE)(**Fig. 1C)**. The backbone atom RMSDs of the best matching pairs of conformers were 0.6 and 0.5 Å (residues 112- 128, 138-152), respectively. In all structures a precisely defined stable hydrophobic core was formed by a Trp154 and several valine, leucine and isoleucine residues between the two α-helices and the N-terminal stretch of irregular secondary structure. Only the middle part of the loop between the helices was less well defined and seemed to have similar loop dynamics. The same residues (Cys132, Arg133, Ala134 or the corresponding Cys, Lys, Gly in C-MANF) had missing amide signals from the ^1^H, ^15^N HSQC spectra, which was an indication of millisecond time-scale chemical exchange (Latge et al. 2015). Also, the low number of NOE peaks arising from atoms in this region suggested flexibility. Thus, cysteines 132 and 135 were disulfide bonded and formed a CXXC motif, but apparently do not tightly constrain conformational dynamics in the loop.

### Biological activity of the C-terminal domain of CDNF

We first tested the neuroprotective effects of GDNF, CDNF, C-CDNF, and N-CDNF in cultured mouse DA neurons where the neuronal cell death was triggered by growth factor deprivation. Consistent with prior studies, GDNF protected mouse DA neurons from cell death, whereas CDNF did not have neuroprotective activity. **(Fig. 2A)**. N-CDNF had no activity. However, surprisingly C-CDNF was active even at 10 ng/ml **(Fig. 2A**). Because CDNF and MANF are neuroprotective when microinjected into neurons, but not when given extracellularly in the model of growth factor deprivation-induced apoptosis (Hellman et al. 2011; Eesmaa et al. 2022), these data strongly suggested that C-CDNF can enter cells. Since C-CDNF production in the CHO cells gave suboptimal yields, we tested the biological activity of chemically synthesized C-CDNF. Remarkably, CHO-derived C-CDNF and chemically synthesized C-CDNF rescued mouse DA neurons in two different *in vitro* models of PD – in an oxidative stress model induced by 6-hydroxydopamine (6-OHDA) and in a thapsigargin triggered ER stress model. In these two models, both C-CDNF preparations demonstrated neuroprotection at the tested 100 ng/ml concentration and their activities were comparable to full-length CDNF and GDNF (**Fig. 2B,C).**

**Figure 2.**
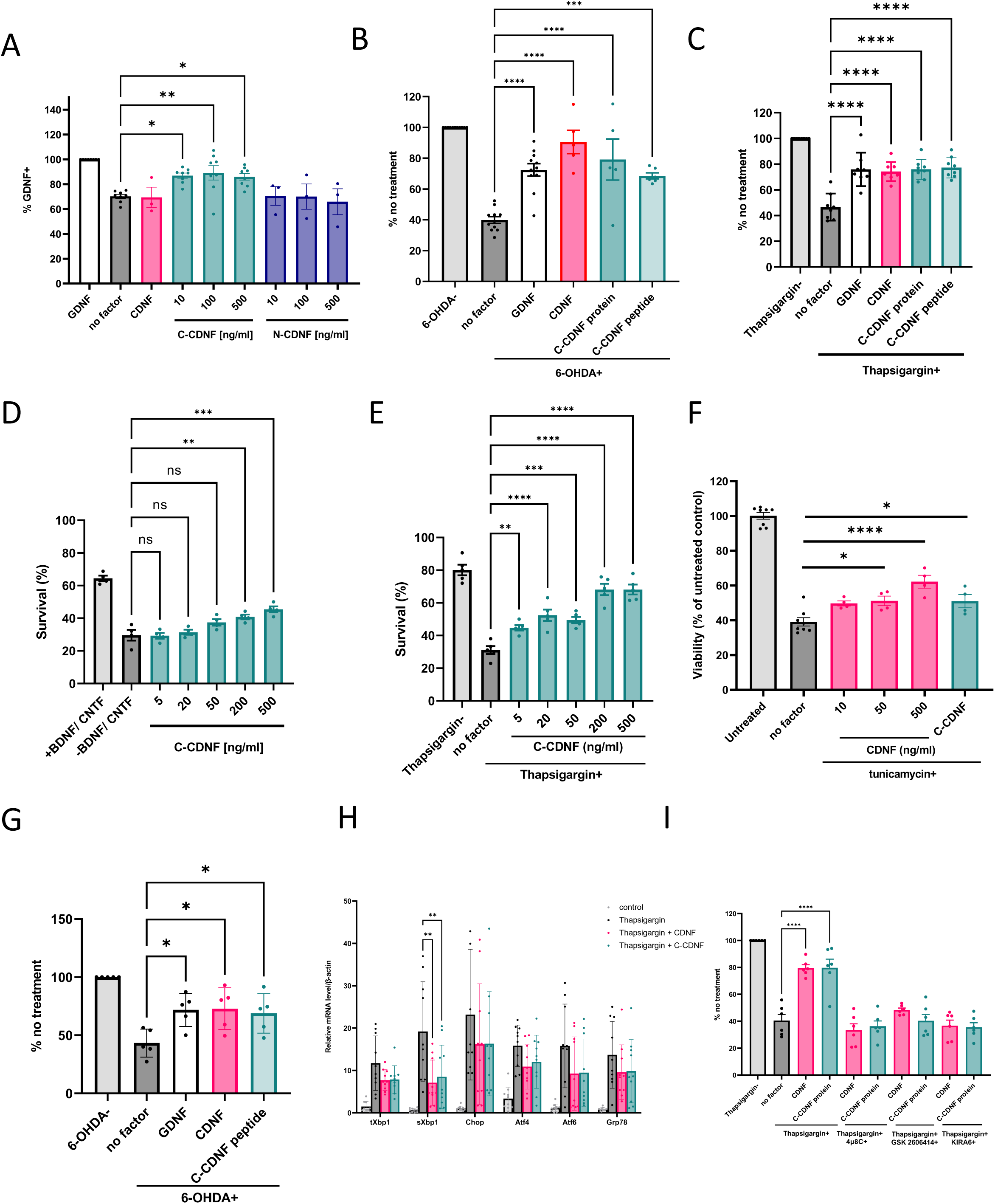
Effect of C-CDNF on the survival of mouse DA neurons and MNs. **A)** C-CDNF increased the survival of mouse dopamine (DA) neurons from growth factor deprivation in a dose- dependent manner. Mouse midbrain DA neurons were cultured with or without recombinant protein C-CDNF or N-CDNF (10-500 ng/ml) for 5 days. DA neurons were identified by TH immunostaining and expressed as % of cell survival in each condition compared to GDNF-treated neurons. Shown are the means ± SEM of 3-8 independent experiments per condition. **B-C)** C-CDNF can protect mouse DA neurons against 6-OHDA or thapsigargin-induced toxicity. DA neurons were treated with **B)** 15 µM 6-OHDA or **C)** 20 nM thapsigargin only, or toxin with the indicated growth factors. After three days, TH+ neurons were counted and expressed as % of cell survival in each condition compared to non-toxin treated neurons. Shown are the means of 5-8 experiments ± SEM **D)** C-CDNF increased the survival of MNs in a dose-dependent manner. MNs cultured 5 days in the presence of different concentrations of C-CDNF (5-500 ng/ml) BDNF/CNTF was used as a positive control. Mean ± SEM of 4 different experiments. **E)** C-CDNF protected MNs against thapsigargin induced toxicity in a dose-dependent manner. MNs were treated with thapsigargin (5 nM) for 24 h in the presence of different dose of C-CDNF (5-500 ng/ml). Mean ± SEM of 5 different experiments. **F)** C-CDNF 50 ng/ml rescued iPS cell-derived MNs from tunicamycin induced cell death. Mean ± SEM of 4-8 t experiments. **G)** C-CDNF protected commercially available iPSC-derived DA neurons against 6- OHDA induced toxicity. **H)** C-CDNF added to the cultured DA neurons decreased the expression levels of UPR transcripts upon 24 h thapsigargin (100 nM) induced ER stress as measured using qPCR (normalization to β-actin). Mean ± SEM of 11 experiments. **I)** Protective effect of C-CDNF against ER-stress is dependent on active IRE1α and PERK pathways. DA neurons were cultured with C-CDNF alone or in combination with 4µ8C, KIRA6 or GSK2606414 and with the treatment of thapsigargin (20 nM) for three days. TH-positive neurons were counted and expressed as % of non- toxin treated neurons. Shown are the means of 6 experiments ± SEM. *P < 0.05, **P < 0.01, ***P < 0.001; one-way ANOVA followed by Sidak post hoc test in A–I.

We previously demonstrated that CDNF rescues MNs *in vitro* and *in vivo* in three distinct rodent models of ALS by primarily inhibiting ER stress-associated cell death (De Lorenzo et al. 2023). Developing MNs depend on trophic support of BDNF and ciliary neurotrophic factor (CNTF) for survival *in vitro.* Similarly to CDNF, C-CDNF had a clear but modest survival-promoting effect on developing MNs compared to BDNF/CNTF in the growth factor deprivation assay **(Fig. 2D).** In contrast, when thapsigargin-induced ER stress was triggered in MNs, C-CDNF rescued MNs from cell death in a dose-dependent manner (**Fig. 2E**). The robust effect of C-CDNF was apparent already at 5 ng/ml and matched our earlier results obtained in DA neurons **(Fig. 2B, C)**.

As a crucial step towards clinical translation, we tested and confirmed the survival-promoting effects of CDNF and C-CDNF on human iPSC-derived MNs and DA neurons. We induced cell death in human iPSC-derived MNs by tunicamycin treatment, which blocks N-glycosylation of proteins and triggers ER stress-induced cell death. Both CDNF and C-CDNF rescued human MNs from tunicamycin-induced cell death at 50 ng/ml **(Fig. 2F)**. In addition, we tested whether C-CDNF could rescue human iPSC-derived DA neurons from 6-OHDA-induced cell death **(Fig. 2G).** Similarly to CDNF and GDNF, C-CDNF rescued human DA neurons. These data constitute the first demonstration that C-CDNF and CDNF protect human MNs and human DA neurons.

We previously found that CDNF efficiently protects peripheral sympathetic superior cervical ganglion (SCG) neurons from tunicamycin induced cell death (Eesmaa et al. 2022). We delivered plasmids encoding C-CDNF or C-CDNF protein by neuronal microinjection into SCG neurons and found that both plasmid encoding C-CDNF or C-CDNF protein rescued SCG neurons from tunicamycin-induced apoptosis. The neuroprotective activity of C-CDNF was comparable to CDNF **(Suppl. Fig. 1A, B).** Taken together, these results clearly demonstrate that similarly to CDNF, its C-terminal domain can protect and rescue CNS DA neurons, MNs and peripheral sympathetic neurons.

### C-CDNF reduces the level of *sXbp1* mRNA and its neuroprotective activity in cultured dopamine neurons is blocked by IRE1**α** and PERK inhibitors

We previously found that MANF and CDNF regulate the UPR by attenuating IRE1α, PERK and ATF6 signalling pathways (Apostolou et al; Arancibia et al. 2018; Lindahl et al. 2014; Voutilainen et al. 2017; Eesmaa et al. 2021; Eesmaa et al. 2022; De Lorenzo et al., 2023; Kovaleva et al., 2023). The use of specific IRE1α and PERK inhibitors demonstrated that the antiapoptotic effect of CDNF in neurons relies on the activities of IRE1α and PERK signaling pathways (Eesmaa et al. 2022; De Lorenzo et al., 2023). Given these prior findings, we tested which mechanisms govern C-CDNF’s favorable effects in protecting neurons from cell death.

C-CDNF rescued mouse embryonic DA neurons from ER stress-induced apoptosis primarily through IRE1α-mediated UPR signaling. We first investigated the expression of key marker genes associated with all three UPR pathways in cultured primary DA neurons. Transcription of spliced *Xbp1* (*sXbp1*) corresponds to the activation of IRE1α pathway. *Xbp1 was* upregulated by thapsigargin and was strongly down-regulated by C-CDNF (**Fig. 2H**). The activation of the PERK pathway was monitored by following expression levels of *Atf4* and *Chop* transcription factors. *Atf4* and *Chop* were upregulated by thapsigargin, and we observed a trend in their down-regulation by C-CDNF. As expected, *Atf6* and its downstream target *Grp78* of the ATF6 pathway were upregulated by thapsigargin, and there was a trend in their down-regulation following C-CDNF **(Fig. 2H).** Taken together, these data suggest that extracellularly applied C-CDNF rescues mouse embryonic DA neurons from ER stress-induced apoptosis by primarily regulating IRE1α mediated UPR signaling. C-CDNF appeared to also regulate PERK and ATF6 branches of the UPR, but these data did not reach statistical significance.

Additional mechanistic studies confirmed that the neuroprotective effects of C-CDNF, similarly to CDNF, depended on UPR signaling from both PERK and IRE1α pathways. We first tested the neuroprotective effects of C-CDNF and CDNF in thapsigargin-treated DA neurons in the presence of PERK kinase inhibitor GSK2606414 (Axten et al. 2012), IRE1α RNase inhibitor 4μ8C (Cross et al. 2012), or IRE1α kinase inhibitor KIRA6 (Ghosh et al. 2014). C-CDNF and CDNF protected DA neurons from thapsigargin-induced cell death (**Fig. 2I**), but their neuroprotective activities were inhibited by PERK and IRE1α signaling inhibitors. We observed similar results with C-CDNF overexpression by microinjection of C-CDNF-encoding plasmid into SCG neurons **(Suppl Fig. 1C).** Collectively, these data demonstrate that the survival-promoting and ER stress-regulating mechanisms of C-CDNF and CDNF are very similar.

### C-CDNF penetrates the cell membranes and reaches the brain and spinal cord

Having established the ability of C-CDNF to protect both DA neurons and MNs *in vitro*, we found that C-CDNF was a cell membrane-penetrating protein. Mouse embryonic day 14 DA neuron cultures were incubated with ^125^I-C-CDNF or ^125^I-CDNF in culture medium. To establish membrane permeability, cells were washed to remove the treated culture medium followed by acid wash to remove proteins bound to the cells surface, then lysed to release membrane-bound and internalized protein linked Iodine-125 radioisotope, which was quantified using a gamma counter. We found that the C-CDNF was enriched in the membrane-bound and internalized cell fractions, in contrast to CDNF **(Fig. 3A)**.

**Figure 3.**
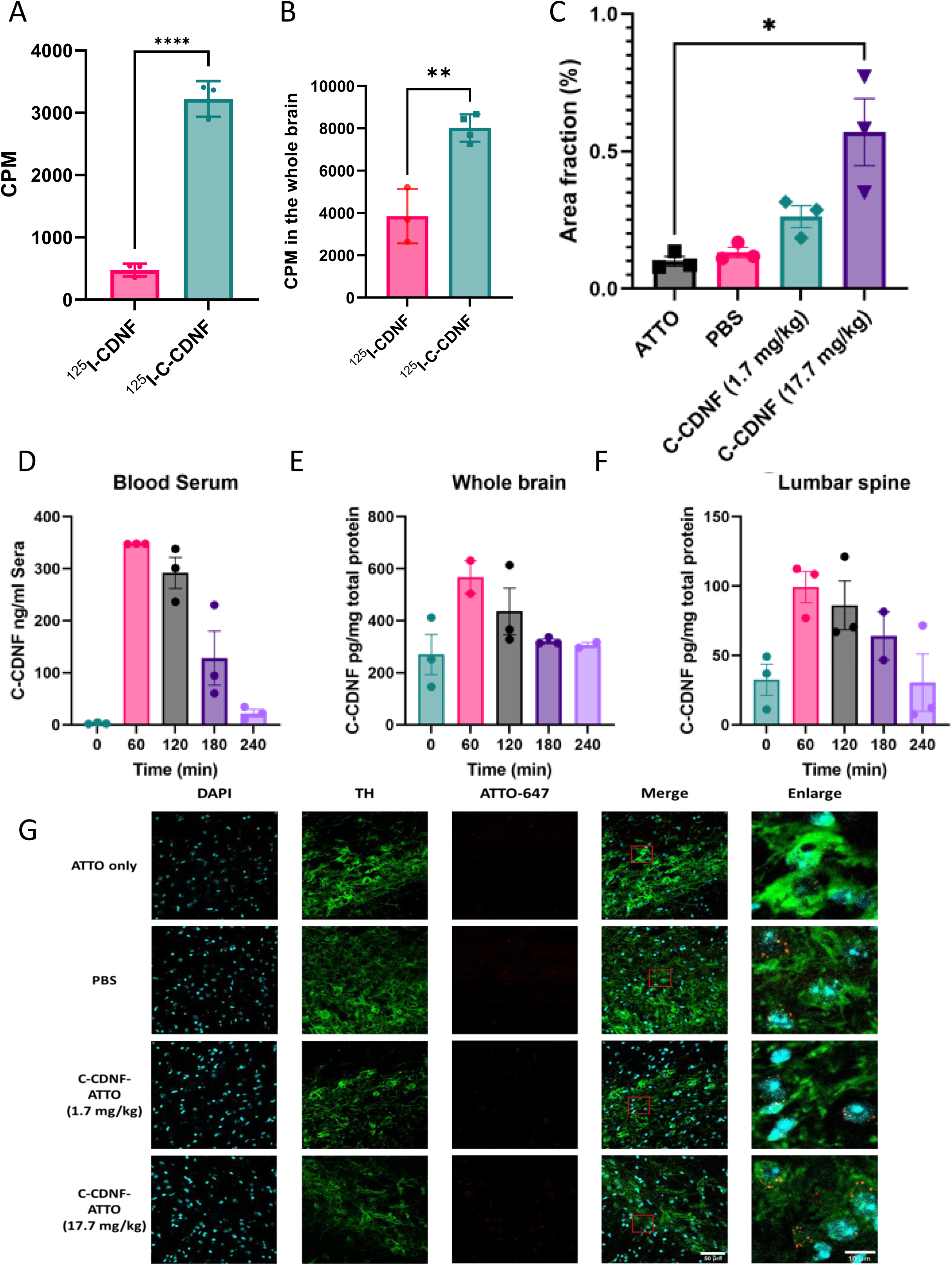
C-CDNF penetrates the cellular membrane and is detected in the brain when injected subcutaneously. **A**) ^125^I-C-CDNF is found in significant concentration from the intracellular fraction (p < 0.0001) after incubation in the media. **B**) After subcutaneous injection, ^125^I-C-CDNF is detected in the rat brain at significantly higher concentration than ^125^I-CDNF (p = 0.0023). **C**) After subcutaneous injection, C-CDNF-ATTO-647 is detected in the SNpc of the rat brain at significantly higher concentration than ATTO-647 vehicle (P = 0.0279). **D-F**) Subcutaneously injected C-CDNF is detected within 60 minutes of injection within the mouse brain and lumbar spinal region as detected by ELISA. **G)** Representative images of subcutaneously injected ATTO-647-linked polypeptides, ATTO-unlinked and PBS.

One key limitation of using full-length CDNF as a therapeutic protein in neurological diseases is that it fails to cross the BBB and thus does not reach target tissues. Consequently, current pre-clinical studies using full-length CDNF require delivery of the therapeutic protein directly to the target site inside the brain, necessitating invasive brain or intrathecal delivery approaches.

Following our discovery that C-CDNF, unlike full-length CDNF, can penetrate the cell membrane, we set out to establish the brain-penetrating properties of C-CDNF. In a first approach, rats were injected subcutaneously with ^125^I-C-CDNF or ^125^I-CDNF. Two hours post-injection, rats were transcardially perfused to clear the brain of blood. With the gamma counter we measured the presence of the ^125^I-labelled CDNF polypeptides present in the brain. We found that C-CDNF was detected at significantly higher concentrations in the brain compared to CDNF indicating that more C-CDNF was transported to the brain **(Fig. 3B).**

In a second approach we labelled C-CDNF with fluorescent compound ATTO-647. In control experiments, we confirmed that C-CDNF-ATTO-647 retained its biological activity **(Supplementary** Fig. 1). We then subcutaneously injected mice with PBS, PBS ATTO-647, or C-CDNF-ATTO-647.

Mice were transcardially perfused to clear the brain of blood two hours post-injection. Using immunofluorescence we quantified the presence of C-CDNF-ATTO in the substantia nigra pars compacta (SNpc) of the mouse brain, a target area of critical importance for PD. Mice injected with high dose (17.7 mg/kg) C-CDNF-ATTO-647 had a robust ATTO signal in the SNpc area, compared to ATTO-647 injected mice. **(Fig. 3C).** In a third and final approach, we assessed the pharmacokinetic properties of C-CDNF. We subcutaneously injected 5 groups of mice with 100 µg of C-CDNF and resected the brain and spinal cord at 60-, 120-, 180- and 240-minute time points following injection. A control, time point (t=0) was injected with PBS. Mice were sacrificed via cardiac perfusion using ice-cold PBS only, and then snap frozen. Following homogenization and total protein quantification, we developed an in-house ELISA to quantify C-CDNF concentrations in the blood serum, lumbar spinal cord, and the left hemisphere of the brain. We determined that the peak concentration of C-CDNF in blood serum was 347 ng/ml at 60 minutes following injection **(Fig. 3D)**, with a half-life of 79.9 minutes. We found that C-CDNF reached the brain (567.7 pg/mg) and spinal cord 99.3 pg/mg) peaking at 60 minutes following peripheral delivery **(Fig. 3E-F).** C-CDNF had a half-life of 57.4 minutes in the brain, and 98.6 minutes in the spinal cord.

### Intrastriatally injected C-CDNF prevents dopamine neuron loss and reduces XBP1s expression in microglia in the 6-OHDA rat model of Parkinson’s disease

Having established the neuroprotective effect of C-CDNF *in vitro*, we discovered that C-CDNF protects and restores nigral DA neurons in 6-OHDA-lesioned rats when delivered directly into the brain. In this model, a unilateral lesion was induced by injecting 6-OHDA into the striatum. Two weeks after 6-OHDA injection, amphetamine-induced rotational behavior was measured to estimate the severity of the lesion **(Fig. 4A**). Rats were divided into treatment groups based on their amphetamine-induced ipsilateral turns at week 2, and C-CDNF or vehicle was then unilaterally injected to the striatum. Ipsilateral turns increased in the vehicle-treated group over time, suggesting that the lesion progressed in rats that received the vehicle control. In contrast, ipsilateral turns did not increase in rats that received C-CDNF **(Fig. 4B)**. When overall rotations were analyzed using the area under curve method, C-CDNF-treated rats rotated less than vehicle-treated rats (**Fig. 4C**). Next, we quantified tyrosine hydroxylase (TH)-positive cells in the SNpc. The number of TH-positive cells on the lesioned side, relative to the intact side, was 22 % in the vehicle treated group, compared to 41 % in the C-CDNF-treated group, clearly demonstrating that C-CDNF prevented neuronal loss in the SNpc (**Fig. 4D)**.

**Figure 4.**
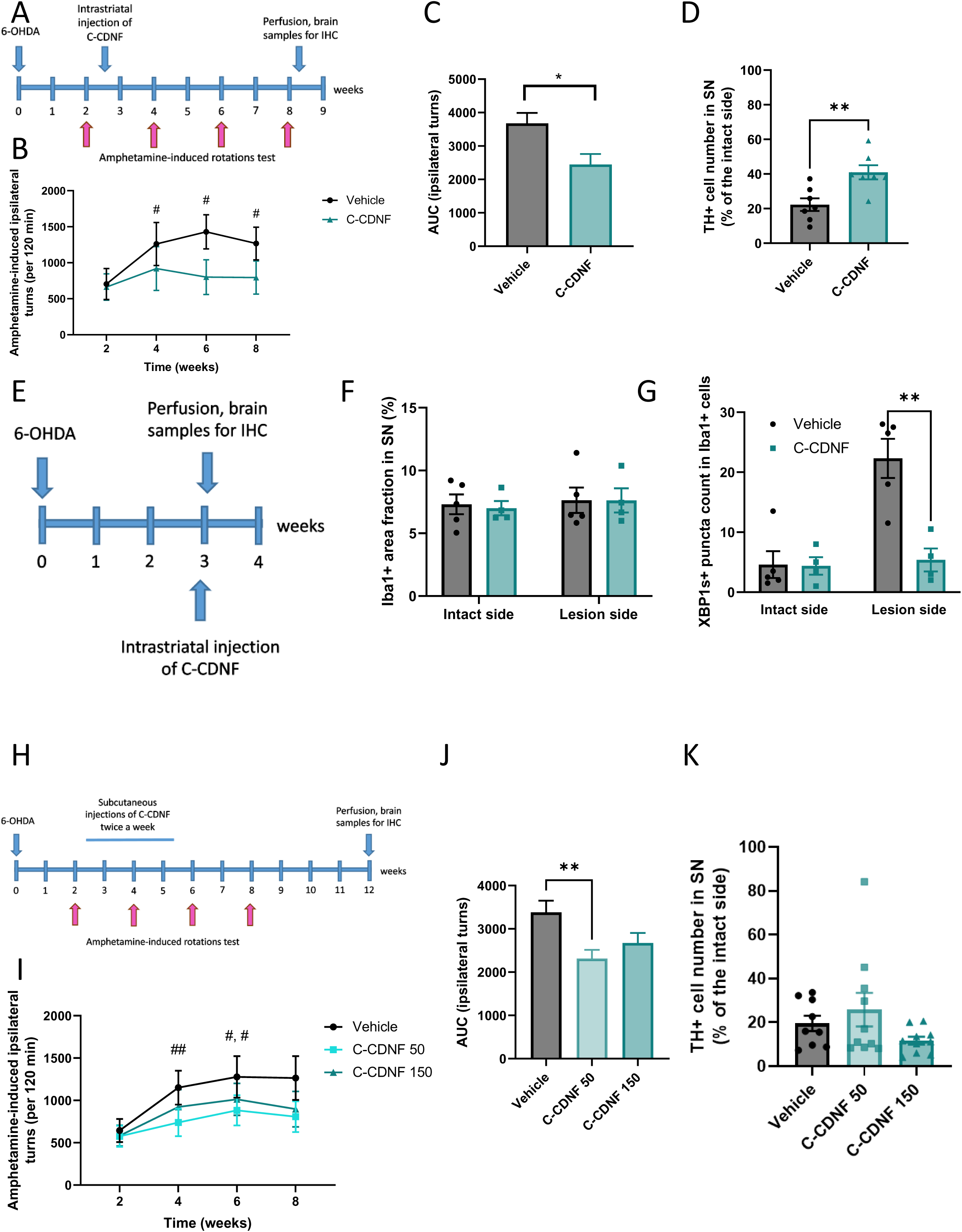
Effects of C-CDNF in a 6-OHDA rat model of Parkinson’s disease. **A)** Timeline of experimental procedure. **B)** Within vehicle group, amphetamine-induced ipsilateral turns were increased at weeks 4, 6 and 8 compared to week 2 (#p = 0.0131, 0.0104 and 0.0393, respectively, repeated measures two-way ANOVA with Dunnett’s multiple comparisons test). **C)** C-CDNF reduced area under curve of amphetamine-induced ipsilateral turns (p = 0.0164, unpaired t test, n = 7). **D)** The number of TH-positive cells in the substantia nigra (SN) (p = 0.0054, unpaired t test, n = 7). **E)** Timeline of experimental procedure. Rats were sacrificed 24 hours after intrastriatal C-CDNF injection. **F)** Iba1-positive cell area in the SN **G)** Number of XBP1s aggregates in Iba1-positive cells in the SN (p = 0.0042, unpaired t test, n = 4–5) **H)** Timeline of experimental procedure. Vehicle, C-CDNF (50 µg per injection) or C-CDNF (150 µg per injection) were injected twice a week for three weeks. **I)** Amphetamine-induced ipsilateral turns (n = 14–17). Within vehicle treated rats, ipsilateral turns were increased from week 2 to week 4 (##p = 0.0086), and to week 6 (#p = 0.0351) (Repeated measures two-way ANOVA with Dunnett’s multiple comparisons test). Within C-CDNF 150 group, ipsilateral turns were increased at week 6 compared to week 2 (#p = 0.0446). **J)** Ipsilateral turns measured using area under curve were significantly decreased in the C-CDNF 50 group compared to the vehicle group (p = 0.007, respectively, one-way ANOVA with Tukey’s multiple comparisons test). **K)** The number of TH-positive cells in the SN (n = 10–11). * p<0.05, ** p<0.01, *** p<0.001 between groups. # p<0.05, ## p<0.01 within group compared to week 2. Results are shown as mean ± SEM.

Next, we evaluated whether C-CDNF had short-term effects on the UPR in Iba1-immunoreactive microglia in 6-OHDA-lesioned rats **(Fig. 4E)**. C-CDNF or vehicle was intrastriatally injected 3 weeks following 6-OHDA injection and rats were sacrificed 24 hours later. We found increased XBP1s- positive aggregates in Iba1-positive cells following 6-OHDA injection, which was reduced by C-CDNF **(Fig. 4G**). The area of Iba1 positivity in the SN was not changed on the 6-OHDA-lesioned brain side compared to the intact side, and C-CDNF did not influence the Iba1-positive cell area (**Fig. 4F).**

Remarkably, C-CDNF restored motor imbalance when peripherally delivered to PD rats. Vehicle or C-CDNF (50 µg or 150 µg) was subcutaneously injected twice weekly for three weeks to 6-OHDA-lesioned rats **(Fig. 4H)**. As seen earlier, amphetamine-induced ipsilateral turns increased over time in the vehicle-treated group. In contrast, no increase in ipsilateral turns was observed in the group that received 50 µg C-CDNF **(Fig. 4I**). When the area under curve was measured, rats treated with 50 µg of C-CDNF, but not 150 µg C-CDNF, rotated less compared to the vehicle-treated group (**Fig. 4J),** demonstrating that lower dose of C-CDNF had an overall protective effect. However, no differences were found in numbers of TH-positive cells in the SN between experimental groups (**Fig. 4K**).

### Weekly subcutaneous C-CDNF injection delays disease onset, preserves motor ability, protects lower motor neurons and reduces microglial activation in a SOD1-G93A mouse model of ALS

Single, weekly peripheral injections of C-CDNF protected MNs from SOD1-associated degeneration and motor function decline in an ALS mouse model. The transgenic (TG) SOD1-G93A mouse model of ALS exhibits a progressive increase in ER stress and UPR activation during course of disease (Kikuchi et al. 2006). Beginning at week 13 (early-stage disease), SOD1-TG mice exhibit a progressive loss of motor ability and physical condition, leading to severe weight loss and eventual total paralysis (until their humane endpoint at approximately 21 weeks of age). We previously showed that a single I.C.V. injection of full-length CDNF delays disease onset and improves motor function (De Lorenzo et al. 2023). Transgenic mice were split into two groups with mixed gender and matched motor ability. At week 13, both TG and wild-type (WT) mice were grouped and received either PBS vehicle (5 ml/kg) or C-CDNF (4 mg/kg). Behavioural tests were performed weekly to measure disease progression, after which mice received a subcutaneous injection of C-CDNF or vehicle until clinical end point (defined as a clinical score of 2) **(Fig. 5A)**. Mice treated with weekly subcutaneous C-CDNF had a slower decline in clinical score from week 14 to week 16 when compared to vehicle-treated SOD1-G93A TG littermates **(Fig. 5A)**. To assess the decline in motor coordination, mice were trained to run on the accelerating rotarod apparatus and assessed weekly. TG mice treated with C-CDNF showed a delay in motor degeneration with a peak difference at week 15 between C-CDNF- and vehicle-treated groups **(Fig. 5B)**.

**Figure 5.**
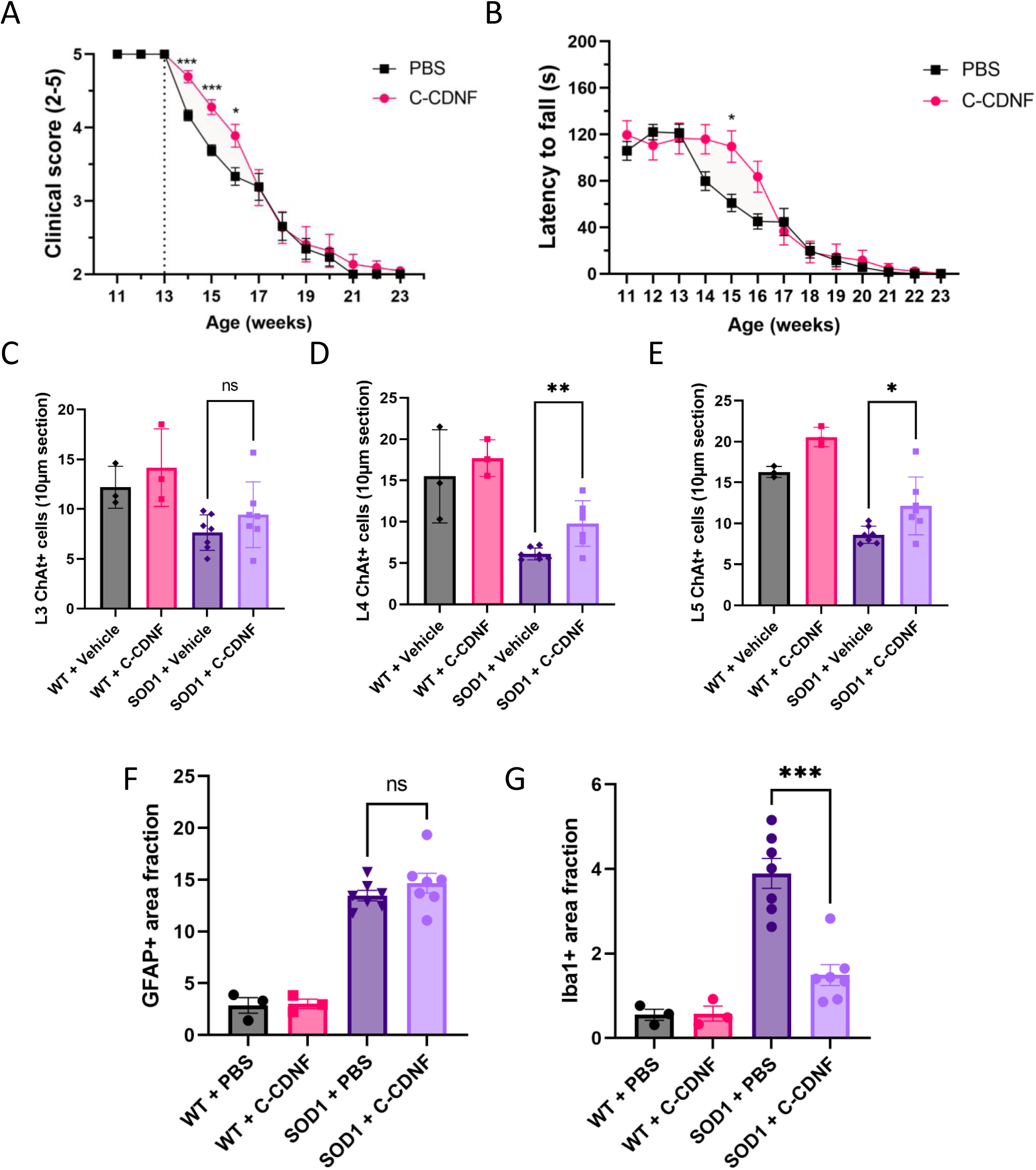
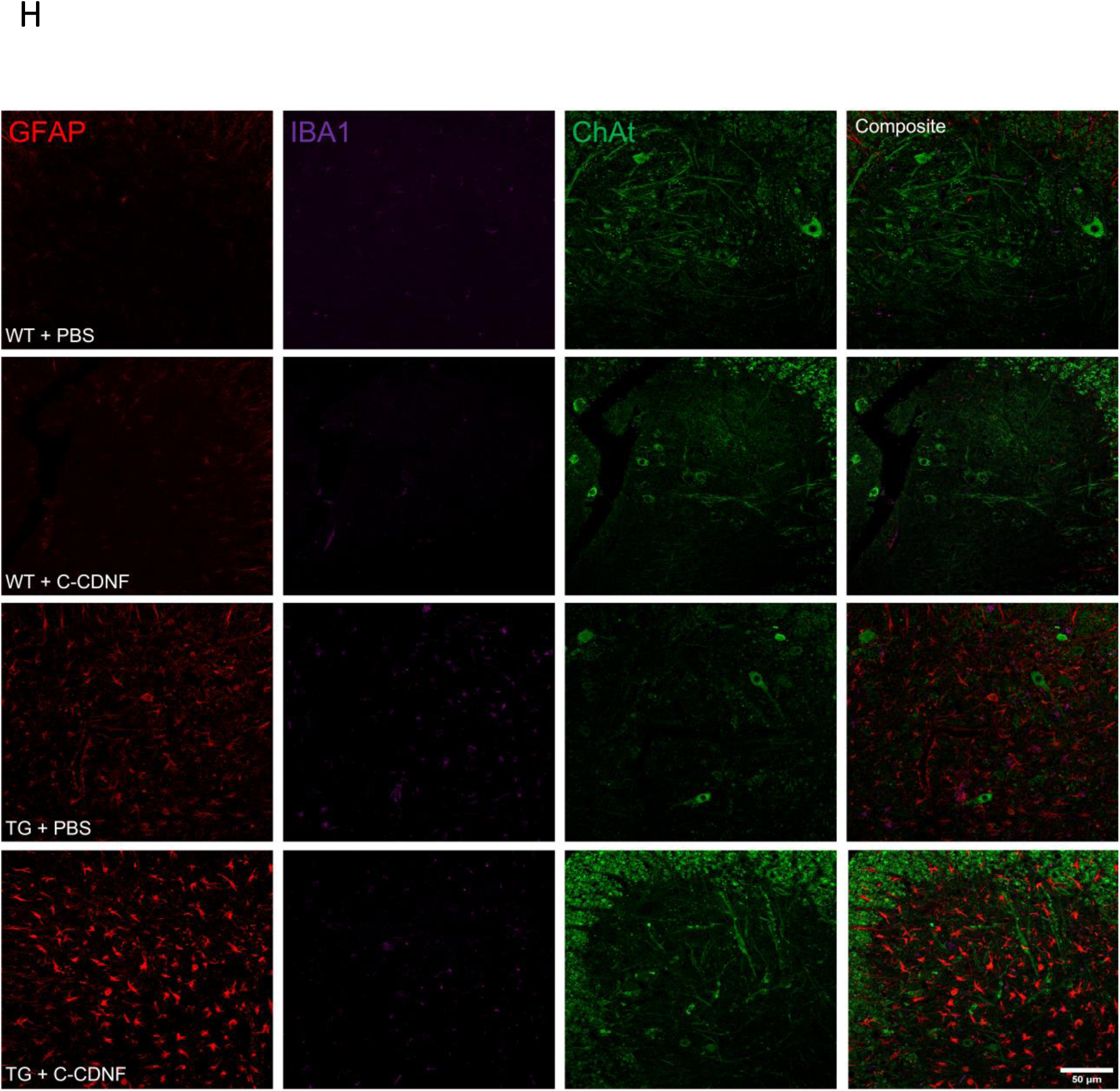
C-CDNF preserves early motor function, protects MNs from degeneration, and reduces microglial activation. **A**) Clinical score of SOD1-G93A mice after weekly injection of C-CDNF (weeks 11-16 n = 18; weeks 17-23 n = 11) or Vehicle (weeks 11-16 n = 21; weeks 17-23 n = 13)showing significant protection at weeks 14 (p = 0.0005), 15 (p = 0.0003), and 16 (p = 0.0402). **B**) Motor ability of mice as tested by weekly rotarod; mice have significant protection at week 15 (p = 0.0331). **C-E**) Immunohistochemistry for MN markers from lumbar area L3-5 with significant protection in C-CDNF (n=7) treated transgenic (TG) mouse MNs versus vehicle (n = 7) treated mice MN count in L4 (p = 0.005) and L5 (p = 0.0257). WT vehicle (n = 3) treated and C-CDNF (n = 3) treated mice MN count for comparison. **F)** Immunofluorescence for markers of astrocytic activation **G)** Immunofluorescence for markers of microglial activation, C-CDNF (n=7) treated TG mice have significantly less microglial activation (p = 0.0001) versus vehicle (n = 7) treated mice, WT vehicle (n = 3) treated and C-CDNF (n = 3) treated mice MN count for comparison **H)** representative images of ChAT-stained lower motor area.

Another group of mice were balanced on motor ability and grouped in C-CDNF or vehicle-treated groups as above with both TG and WT controls **(Fig. 5A)**. These groups were treated with single weekly subcutaneous injections of C-CDNF and subjected to weekly behavioural assessments. These cohorts of mice were sacrificed at week 16 to assess the progression of cellular pathology in the lumbar spinal cord. We quantified the number of choline acetyltransferase (ChAT)-positive cell bodies in the ventral horn of the lumbar spinal cord (L3 to L5). We discovered a higher number of ChAT positive MNs in L4 and L5 in the C-CDNF-treated group compared to vehicle controls at week 16 **(Fig. 5C-E)**. Microglial and astrocytic changes develop as part of the SOD1-linked disease pathology (Nagai et al. 2007). Remarkably, C-CDNF-treated mice had a significantly lower microglial activation state, as measured by Iba1-positive area fraction in the ventral horn of the L5 lumbar spine **(FIG. 5G)**. We did not observe any effect on astrocytic activation, evaluated by GFAP positive signal in the ventral horn of the L5 lumbar spinal cord **(Fig. 5F)**. Notably, in control experiments, we did not detect any microglial or astrocytic changes in wild type mice treated with C-CDNF.

## Discussion

GDNF, neurturin (NRTN) and Platelet Derived Growth Factor Subunit B (PDGF-B) have been tested in animal models of PD and in moderate neurotoxin lesion models: when 30-75% of the nigrostriatal DA neurons were lost, all three factors protected DA neurons and significancy alleviated the motor behavior impairment of lesioned animals (Sidorova and Saarma, 2020). Although according to post hoc analysis of the previous phase 2 clinical trial in PD, GDNF showed therapeutic effects in about 43% of patients (Whone et al. 2019), overall, the phase II clinical trial results with GDNF, NRTN and PDGF-B have been inconclusive (Manfredsson et al. 2020; Barker et al. 2020; Sidorova and Saarma, 2020). Why has the delivery of these trophic factors to the brain not shown therapeutic efficacy? One of the reasons is the intracranial delivery itself. As brain delivery of growth factors is an invasive and risky procedure, ethical committees have only allowed the treatment of later-stage patients with 25% or less of nigrostriatal neurons remaining. The treatment is thus started when most of DA neurons and their projections to the caudate putamen have been lost. Secondly, a very significant problem is the poor diffusion of the administered proteins in the brain parenchyma.

According to published data, significantly less than 50% of the caudate putamen was covered by the growth factors, meaning that more than a half of nigrostriatal DA neurons never met delivered growth factors (Sidorova and Saarma, 2020). Finally, the local delivery of trophic factors with intracranial catheters results in a very high local concentration, that is not optimal for neuroprotection. Therefore, it is important to develop novel approaches to overcome these hurdles. One of the possibilities is to search for growth factors that have better pharmacokinetic properties and the ability to pass through the BBB. During CDNF purification we noticed that some portion of the recombinant CDNF protein is cleaved in the hinge region yielding the N-terminal domain (Parkash et al. 2009). Based on that we predicted that the C-terminal domain (C-CDNF) is also formed. The biological activity of extracellular N-CDNF and C-CDNF was tested in DA neurons in a neuronal cell death model induced by growth factor deprivation. Only C-CDNF was able to rescue DA neurons. This was surprising, as full-length CDNF was also earlier shown to be inactive in this type of assay (Lindholm et al. 2007; Eesmaa et al. 2022). Since microinjected CDNF rescues growth factor deprived neurons via intracellular mechanisms, we postulated that C-CDNF might penetrate the cell membrane. In the following experiments we found that C-CDNF, and not full length CDNF, efficiently penetrates the plasma membrane of DA neurons and rescues them from 6-OHDA- and ER stressor thapsigargin-induced cell death in culture conditions. It is important to note that C-CDNF protects and rescues DA neurons as efficiently as CDNF and is also neuroprotective to MNs. In the presence of BDNF and CNTF, MNs survived, but depriving these neurons from growth factors triggered neuronal cell death. CNTF-BDNF deprivation induced MN death was partially blocked only at high concentrations of C-CDNF. Similar to full-length CDNF, MN cell death induced by the ER stressor thapsigargin can be almost completely blocked by C-CDNF even at very low concentrations. These experiments demonstrate that C-CDNF recapitulated one of the unique properties of CDNF – the ability to rescue ER stressed or 6-OHDA-injured neurons, while having minimal neuroprotective effects on naïve neurons. This property of C-CDNF would be very valuable in clinical trials, reducing potential side effects.

As an important step towards clinical translation, we tested the ability of C-CDNF to protect and rescue cultured human DA neurons and MNs. Since data on the survival promoting effects of CDNF on human DA neurons and MNs are missing, we first tested the effect of full-length CDNF. Results shown in Fig. 2 clearly demonstrate that CDNF efficiently protected human DA neurons from 6- OHDA treatment-induced cell death. Importantly, C-CDNF at the tested concentration of 100 ng/ml also protected human DA neurons.

CDNF protects mouse MNs from ER stress-induced cell death and potently stimulates their neurite outgrowth (DeLorenzo et al. 2023). As no data about the effects of CDNF on human MNs have been published, we first found that C-CDNF at 50 ng/ml and 500 ng/ml protected MNs from tunicamycin-induced cell death. Moreover, C-CDNF protected human iPSC-derived MNs at 50 ng/ml as potently as CDNF, demonstrating that C-CDNF protects human MNs as well.

Earlier data have clearly demonstrated that CDNF can regulate UPR pathways and reduce aggregation of α-synuclein both *in vitro* and *in vivo* models of PD (Voutilainen et al. 2017; Albert et al. 2021; Eesmaa et al. 2021; DeLorenzo et al. 2023; Eesmaa et al. 2022). Furthermore, inhibiting PERK and IRE1α with pathway-specific inhibitors demonstrated that the survival promoting action of CDNF in DA neurons and MNs is inhibited by blocking these pathways (Eesmaa et al. 2022; De Lorenzo et al. 2023). To test whether C-CDNF also regulates UPR pathways, we analyzed the mRNA levels of GRP78, tXbp1, Xbp1s (IRE1α pathway), Chop and Atf4 of PERK pathways and Atf6. Results presented in Fig. 2H shows that C-CDNF significantly reduces Xbp1s mRNA levels and there is a clear trend in decreasing the levels of mRNAs upregulated in PERK and ATF6 pathway during ER stress in DA neurons. Importantly, PERK pathway inhibitor GSK2606414 or IRE1α kinase inhibitor KIRA6 or RNase inhibitor 4µ8C inhibited CDNF’s ability to promote the survival of DA neurons. These data clearly show that, like full-length CDNF, the survival promoting signaling of C-CDNF requires functional PERK and IRE1α pathways, and that C-CDNF can regulate UPR pathways. C-CDNF regulated the IRE1α pathway most prominently.

CDNF and MANF differ from other NTFs and growth factors by their structure and by their mode of action. As mentioned earlier, CDNF consists of two domains, the saposin-like N-terminal domain and SAP-like C-terminal domain. Based on the crystal structure (Parkash et al. 2009) and solution NMR structure (Latge et al. 2015), both domains have independent structures stabilized with disulfide bridges. In this work, we have determined the 3D structure of the C-terminal fragment of CDNF. Our data show that C-CDNF adopts the same fold as the C-terminal domain of mature CDNF (Latge et al. 2015). It thus establishes an independently folding structural and functional module. In our previous work with the CDNF paralogue MANF, we observed that the C-terminal domain of MANF (C-MANF) also folds autonomously and in the mature MANF, the N-terminal Saposin-like domain and the C-terminal SAP domain show no interaction owing to the flexible linker between the domains (Hellman et al., 2011). Remarkably, the surface charge distributions of C-CDNF and C-MANF slightly differ due to the few amino acid substitutions in their primary sequences (Fig. 1C), meaning that this property is likely to be insignificant for the ability of these domains to pass the BBB. C- MANF loses its intracellular activity if the conserved CXXC motif is mutated (C151S, Mätlik et al. 2015). It would be interesting to test if this mutation also affects the BBB passage of C-MANF or C-CDNF.

Perhaps one of the most exciting findings of this study is the demonstration that C-CDNF crosses through cell membranes and can be detected in mouse and rat brain after peripheral delivery. Presence of subcutaneously injected C-CDNF in the brain and midbrain of rodents was verified with several methods. ^125^I-labelled C-CDNF was used first, and then this finding was confirmed using Atto-647 fluorescence labelled C-CDNF. C-CDNF was detected in the nigrostriatal pathway and DA neurons using immunofluorescence. Finally, we used C-CDNF specific ELISA to quantitatively detect C-CDNF concentration in the brain, in the spinal cord and in the sera of rodents. According to the ELISA measurements about 1-2% of the injected C-CDNF reaches the brain. ELISA analysis also allowed us to measure the stability of the systemically delivered C-CDNF, demonstrating that it possesses a half-life of approximately 55-100 minutes, depending on the model. Considering future clinical trials, it is quite clear that both the efficacy of BBB penetration as well as the half-life of C-CDNF require optimization.

To test the neuroprotective efficacy of C-CDNF, it was stereotactically injected into the striatum of 6-OHDA lesioned rats. Direct intracranial delivery C-CDNF restored motor impairment, as assessed by monitoring amphetamine-induced ipsilateral rotations. C-CDNF was efficiently protecting nigrostriatal DA neurons. When C-CDNF was delivered by peripheral administration to 6-OHDA lesioned rats, motor imbalance was also recovered. However, no statistically significant protection of TH-positive DA neurons was detected. The amount of C-CDNF that reached the midbrain target site after peripheral delivery might have been insufficient to provide neuroprotection, and application of a higher or more frequently systemically administered dose might be necessary to achieve the protection of nigral DA neurons in the 6-OHDA neurorestoration model. On the other hand, 4 mg/kg of C-CDNF once a week for SOD1-G93A mice was able to increase motor coordination and protect MNs in the spinal cord. In our recent study, we showed that CDNF improved motor behavior, protected MNs in lumbar spinal cord and regulated ER stress in three different genetic rodent models of ALS (De Lorenzo et al. 2023). We therefore conclude that in the SOD1-G93A mouse model of ALS, C-CDNF had clear therapeutic effect.

Having shown behavioral effects in a PD model and therapeutic effects of systemically delivered C-CDNF in an ALS model, a question arises about possible side effects. We have shown earlier that CDNF and the related MANF do not act on naïve cells but protect and rescue mostly stressed and injured cells (Airavaara et al. 2012; Voutilainen et al. 2011; Eesmaa et al. 2022; DeLorenzo et al. 2023; Kovaleva et al. 2023). It is logical to assume that this unique mode of action of C-CDNF is beneficial when the drug is systemically delivered either to animal models or to patients. Indeed, here C-CDNF was systemically delivered during a relatively long period with no side effects observed in rodents. Another problem might be that systemically delivered CDNF and C-CDNF can activate the immune system and elicit an immune response. This is indeed a possible scenario but CDNF and possibly also C-CDNF can be detected in the blood of healthy persons meaning that these proteins are not very strong elicitors of the immune response. Furthermore, CDNF was delivered to the patients in clinical trials and no CDNF antibodies were detected (Huttunen et al. 2023). Although we detected clear therapeutic effect in animal model of ALS, in rat 6-OHDA model systemic delivery of C-CDNF had no statistically significant neuroprotective effect on DA neurons. Since we have noticed no neuroprotective effect also when too low doses of CDNF were injected into the brain of 6-OHDA treated rats one plausible explanation is that C-CDNF did not reach the midbrain at sufficient concentrations. Consequently, both the half-life and BBB penetration of the C-CDNF must be improved. University of Helsinki has licensed C-CDNF technology (Saarma, Airavaara, Voutilainen et al. 2018) to Herantis Pharma Plc. and the company has now optimized C-CDNF for clinical trials (Kulesskaya et al. 2024).

The innovative aspect of this study is the new groundbreaking concept for treating neurodegenerative diseases using peripheral delivery of BBB-penetrating CDNF-derived polypeptide with neuroprotective properties, which allow for the avoidance of risky brain surgery. Peripheral delivery also enables disease-modifying treatment to start immediately upon diagnosis of ALS and PD. Earlier medication results in better treatment outcomes since more neurons and neuronal circuits can be rescued and protected from degeneration.

## Methods

### NMR spectroscopy and structure determination

The C-terminal domain of CDNF (residues 101-161) was expressed and purified using the protocol described for MANF (Hellman et al. 2011). The C-terminal domain of CDNF was produced and purified using the protocol described for MANF (Hellman et al. 2011). The NMR sample contained 1 mM C-CDNF in 20 mM sodium phosphate, 50 mM NaCl, 93/7 % H2O/D2O, pH 6.5 buffer. NMR experiments were performed at 25 °C using a Varian INOVA 600 MHz spectrometer equipped with a cryogenically cooled 5 mm {^15^N, ^13^C}^1^H triple-resonance z-gradient probehead and a Varian INOVA 800 MHz spectrometer equipped with a cryogenically cooled 5 mm {^15^N, ^13^C}^1^H triple-resonance z-gradient probehead. Protein backbone chemical shifts were assigned using a suite of 3D triple-resonance experiments, that is, iHNCACB (Tossavainen and Permi 2004), HN(CO)CACB, HNCA, HN(CO)CA, HNCO and HCACO (Permi and Annila 2004). For aliphatic side chain chemical shift assignments 3D H(CCO)NH, (H)C(CO)NH, HCCH-COSY and for aromatic assignments 2D (HB)CB(CGCD)HD, (HB)CB(CGCDCE)HE (reviewed in Sattler et al., 1999) experiments were collected. In addition, the double sensitivity-enhanced DE-HCCmHm-TOCSY experiment was employed to assignment of methyl containing residues (Permi et al. 2004). Aromatic assignments were completed using a NOESY-^13^C-HSQC spectrum. The latter together with a NOESY-^15^N-HSQC spectrum was used to derive distance restraints. Additionally, TALOS-N dihedral angle restraints were derived from chemical shifts (Shen and Bax, 2013). All spectra were processed with NMRPipe (Delaglio et al. 1995) and analysed with CcpNmr Analysis (Vranken et al. 2005) v. 2.5.2. CYANA v. 3.98.15 (Guntert and Buchner 2015) automated structure calculation program was used to generate 300 C-CDNF structures. Of those, 30 with the lowest target function values were further refined in explicit water using AMBER 22 (Case et al, Amber 2022, UCSF). Twenty structures best conforming to given restraints were chosen to represent the structure.

### Midbrain dopamine neuron cultures and C-CDNF treatment

The midbrain floors were dissected from the ventral mesencephalic of 13 days old NMRI strain mouse embryos (E13) following the published procedure (Yu et al 2008). The tissues were incubated with 0.5 % trypsin (103139, MP Biomedical) in HBSS (Ca^2+^/Mg^2+^-free) (14170112, Invitrogen) for 20 min at 37 °C, then mechanically dissociated. Cells were plated onto the 96-well plates coated with poly-L-ornithine (Sigma-Aldrich). Equal volumes of cell suspension were plated onto the center of the dish. The cells were grown for 5 days without any added neurotrophic factor in DMEM/F12 medium containing N-2 supplement (17502048, Invitrogen). For survival assay, the cells were grown with different concentrations of C-CDNF protein or C-CDNF peptide (10 ng/ml-1 µg/ml). Human recombinant GDNF (100 ng/ml) (P-103-100, Icosagen), CDNF protein (100 ng/ml) (P-100-100, Icosagen) or a condition without any trophic factor were used as positive and negative controls, respectively. After growing 5 days, the neuronal cultures were fixed and stained with anti-tyrosine hydroxylase (TH) antibody (MAB318, Millipore Bioscience Research Reagents). Images were acquired by ImageXpress Nano automated imaging system (Molecular Devices, LLC). Immunopositive neurons were counted by CellProfiler software, and the data was analyzed by CellProfiler analyst software (McQuin et al. 2018). The results are expressed as % of cell survival compared to GDNF-maintained neurons (Mahato et al. 2020).

For rescue from neurotoxin treatment, the cells were grown for 5 days without any trophic factors in 96-well plates. Then, the cells were treated with 6-hydroxydopamine hydrochloride (6-OHDA, 15 µM, H4381, Sigma Aldrich) or thapsigargin (20 nM) (T7458, Thermo Fisher Scientific) and C-CDNF protein or C-CDNF peptide (100 ng/ml or 100 nM). After 3 days, the neuronal cultures were fixed and stained with anti-TH antibodies.

### Primary embryonic motoneuron culture and survival assays

Murine embryonic spinal MNs were isolated and cultured as described before (De Lorenzo et al., 2023). Briefly, the spinal cord of E13 embryos was dissected and incubated for 15 min in 0.1 % trypsin (Worthington) in Hank’s balanced salt solution (Thermo Scientific). Subsequently, 0.1 % trypsin inhibitor (Sigma Aldrich) was added to the tissue. Spinal cords were triturated and incubated in Neurobasal medium (Thermo Scientific), supplemented with 1× Glutamax (Thermo Scientific) on Nunclon plates (Nunc) pre-coated with antibodies against the p75 NGF receptor (MLR2, provided by Mary-Louise Rogers and Robert Rush, in collaboration with Flinders University (Adelaide, Australia) for 45 min. Plates were washed three times with Neurobasal medium, and the remaining MNs were recovered using depolarization solution (0.8% NaCl, 35 mM KCl, and 2 mM CaCl2) and collected in MN medium (2% horse serum, 1x B27 in Neurobasal medium with 1x Glutamax).

Isolated MNs were plated at a density of 1,000 cells per cm^2^ on poly-ornithine/laminin-coated dishes. To examine the effect of C-CDNF on the survival of cultured MNs, the cells were counted 4 h after plating to determine the total number of plated cells. MNs were cultured in the presence of the indicated concentrations of C-CDNF and counted again after 5 days in culture to determine the number of surviving cells. As control conditions, MNs were cultured with 10 ng/ml of BDNF and CNTF or without trophic support. To analyze the protective effect of C-CDNF against thapsigargin-induced ER stress, we determined the total number of cells before adding 5 nM thapsigargin. The cells were counted again 24 h after the treatment to determine the number of surviving cells.

### Human iPSC-cell derived motoneurons

hIPSC line HEL24.3 was used for production of hIPSC-derived MNs (Trokovic et al. 2015). hIPSC- derived MNs were produced from hIPSCs according to previously established protocol with minor modifications (Selvaraj et al. 2018). At day 38 of MN differentiation hIPSC-derived MNs were plated at a density of 100 000 cells per cm^2^ on poly-ornithine/laminin/fibronectin/Matrigel-coated dishes. To analyze the protective effect of C-CDNF against tunicamycin-induced ER stress, we washed out the supporting growth factors from the hIPSC-derived MN cultures. After 24h growth factor deprivation period, we treated the MNs with 80 µg/ml tunicamycin and CDNF (10, 50, or 500 ng/ml) or C-CDNF (50 ng/ml) for 4h. After 4h treatment, we washed tunicamycin out from the cultures and maintained MNs with the growth factors for an additional 48h. Cell viability at different conditions was analyzed using PrestoBlue -cell viability assay. Results are presented as the percentage of viable cells compared to untreated control condition. As a negative control condition, MNs were treated with tunicamycin and cultured for 48h without trophic support.

### Human iPS-cell derived dopamine neurons and C-CDNF treatment

iCell® DopaNeurons (01279, FUJIFILM Cellular Dynamics) were seeded according to FUJIFILM Cellular Dynamics user protocol onto the 96-well plates coated with poly-L-ornithine (P3655, Sigma-Aldrich) and laminin (L2020, Sigma-Aldrich). Equal volumes of cell suspension were plated onto the center of the dish. The cells were grown for 5 days in iCell Neural Base Medium 1+ iCell Neural Supplement B (M1010, M1029, FUJIFILM Cellular Dynamics). Then the cells were treated with 6- OHDA (50 µM) and C-CDNF polypeptide (100 nM), without any trophic factor or with GDNF (100 ng/ml) as negative and positive control, respectively. After 3 days, the cells were fixed and stained with anti-TH antibody. Images were acquired by ImageXpress Nano automated imaging system. Immunopositive neurons were counted by CellProfiler software, and the data was analyzed by CellProfiler analyst software. The results are expressed as % of cell survival compared to non-treated neurons.

For the experiments with IRE1α and PERK inhibitors, mouse E13 midbrain dopamine neurons were isolated and cultured for 5–7 days as described and then treated with thapsigargin (20 nM). Recombinant CDNF protein, C-CDNF protein or C-CDNF peptide (100 ng/ml) and IRE1α inhibitors 4μ8C (10 μM) (HY-19707, MedChemExpress), KIRA6 (200nM) (HY-19708, MedChemExpress) or PERK inhibitor GSK2606414 (T2614, argetMol Chemicals) were added to the culture medium at the same time. After 3 days the neuronal cultures were fixed and stained with anti-TH antibodies.

All animal experiments were carried out following European Community guidelines for the use of experimental animals and approved by the Finnish National Experiment Board (License number: ESAVI/12830/2020) and by the Laboratory Animal Center of the University of Helsinki (license no. KEK20-015; 2.7.2020).

### RNA isolation, reverse transcription and quantitative PCR

Midbrain dopamine neurons were cultured for 5-7 days and then treated with thapsigargin (200 nM). Recombinant CDNF protein or C-CDNF (100 ng/ml) was added to the cultures at the same time. After 24 h, RNA from cultured cells was isolated by TriReagent® (RT118, Molecular Research Center) according to manufactureŕs instructions. RNA was reverse transcribed to cDNA with RevertAid™ Premium Reverse Transcriptase (EP0441, Fermentas UAB, Thermo Fisher Scientific Inc). Quantitative PCR was performed using LightCycler® 480 SYBR Green I Master (04887352001, Roche Diagnostics GmbH) and Roche LightCycler® 480 Real-Time PCR System (Roche Diagnostics GmbH). The expression levels were normalized to the levels of β-actin in the same samples. Primers for Grp78, Chop, sXbp1, tXbp1, Atf4, Atf6α used in quantitative PCR were synthetized using previously published sequences (Lindahl et al. 2014; Cao et al. 2016; Danilova et al. 2019; Eesmaa et al. 2021).

### *In vitro* permeability studies

#### Neuronal uptake experiments

Mouse E14 DA neurons (Yu et al. 2003) were cultured for 5 days *in vitro* and incubated with 30 000 cpm of ^125^I-C-CDNFor ^125^I-CDNF at 37°C for 2 h. The cells were transferred to ice and washed once with 0.5 ml of ice-cold culture medium. Then the cells were transferred to Eppendorf tubes and washed at 4 °C once with 0.2 M acetic acid, 0.5M NaCl, pH 2.8. After centrifugation at 1000 g for 10 min the cells were dissolved in 0.5 ml of 0.5 N NaOH and the cell-bound/internalized radioactivity was determined with a gamma counter (Wizard 3, 1480 Automatic Gamma Counter, Wallac/PerkinElmer).

### Experimental animals

Male Wistar rats (weight 220-280 g, at the beginning of the study) and male C57BL/6JRccHsd mice (Envigo, Netherlands) were housed in groups of 3 to 4 under a 12 h light-dark cycle at an ambient temperature of 20–23 °C. Food pellets (Harlan Teklad Global diet, Holland) and tap water were available ad libitum. The experiments were conducted according to the 3R principles of EU directive 2010/63/EU on the care and use of experimental animals, as well as according to local laws and regulations, and were approved by the national Animal Experiment Board of Finland (protocol approval numbers ESAVI/7812/04.10.07/2015 and ESAVI/12830/2020). All experiments were performed in a blinded manner and the animals were assigned to the treatment groups randomly.

### *In vivo* biodistribution studies

#### Injection of iodinated study compounds, tissue dissection and measurement of radioactivity

Specific activity of the iodinated samples in this study was approximately 10^8^ cpm/µg (0.45 µCi/µg). Each animal was injected s.c. with a single 500 000 cpm (∼5 ng) dose, in a 100 µl injection volume. The rats were anesthetized with an overdose of sodium pentobarbital (90 mg/kg, i.p.; Orion Pharma) and perfused transcardially with PBS followed by 4% paraformaldehyde (PFA; Sigma) in a 0.1 M sodium phosphate buffer, pH 7.4. In order to quantify the amount of ^125^I-CDNF and ^125^I-C-CDNF, the brain was removed. BBB crossing ability of the test compounds was assessed 2 h after s.c. injection of radiolabeled CDNF or C-CDNF based on detection of radioactivity (counting with Wizard 3, 1480 Automatic Gamma Counter, Wallac).

#### Injection of Atto-647-C-CDNF, tissue dissection, immunohistochemistry and confocal microscopy

100 µl [1.77 mg/kg (low dose) or 17.7 mg/kg (high dose)] of Atto-647-labeled C-CDNF or Atto-647 dye alone was injected s.c. to naïve mice (n=3 per group). 3 h later the mice were deeply anesthetized using pentobarbital (Orion Pharma, Finland) and thereafter transcardially perfused using cold PBS and 4 % PFA in PBS. After the perfusion, brains were dissected out and post-fixed with 4 % PFA in PBS for 24h at +4 °C. Next day, brains were transferred into 20% sucrose (Sigma) solution. After brains sunk, 35-µm thick coronal sections were cut on a Leica CM3050 cryostat and processed for immunohistochemical staining. First, the sections were mounted on the glass slide and dried. Thereafter, they were rounded using Oil pencil and rinsed with PBS 3 times for 10 min. Next, the tissues were permeabilized and blocked with 0.2% Triton X-100/1 % BSA in PBS for 1 h at room temperature. Then, the sections were incubated with the rabbit anti-tyrosine hydroxylase antibody (Pel-Freez, Cat. no. P40101-150; dilution 1:1000) for overnight at + 4 °C. The next day, the tissues were washed with PBS three times for 10 min and thereafter the sections were incubated with Alexa488-conjugated donkey anti-rabbit IgG (Thermo Fisher/Molecular Probes Cat. no A21206; dilution 1:500) at room temperature for 1 h. The tissues were then rinsed with PBS three times for 10 min. For counterstaining, the nuclei were stained with DAPI (Thermo Fisher scientific, 1:1000, Cat.62247) for 10 min at room temperature. Finally, the tissues were washed with PBS twice for 5 min and mounted. The sections were viewed and imaged using confocal microscopy (Carl Zeiss, Germany; model LSM700). Images (1024x1024 pixels) were acquired using constant laser intensity using LD LCI PlanApochromat 25x/0.8 Imm Korr DIC M27 lens (Carl Zeiss) and the following filters: 405 nm for DAPI, 488 nm for TH, and 639 nm for Atto647. Images were acquired in sequential scanning mode averaging 4 images.

#### Atto-647 labelling of C-CDNF

Atto-647 is a red fluorescent label (excitation: 647 nm/emission: 667 nm) characterized by strong absorption, high fluorescence quantum yield, high photostability and good water solubility. C-CDNF was labelled with Atto-647 NHS ester (Sigma-Aldrich) following the procedure recommended by the manufacturer. Atto-647 NHS ester was dissolved in 0.1 M bicarbonate buffer, pH 8.3, and mixed with the protein in a ratio 5:1. The mixture was incubated at room temperature for 1 h and purified by gel filtration. At the pH 8- 9 range, the ε-amino groups of lysines are unprotonated to allow fast coupling, and the Atto-647 NHS-ester readily reacts with ε-amino groups of proteins/peptides. C-CDNF has 7 lysines available for NHS ester-mediated coupling. The biological activity of the Atto- 647-labeled C-CDNF was studied using microinjection to mouse SCG sympathetic neurons and testing their neuronal survival-promoting activity. SCG neurons were prepared from postnatal day-old mice (Yu et al. 2003), cultured for 7 days, and then microinjected with Atto-647-C-CDNF. Next day 2 µM tunicamycin, an inducer of ER stress, was added, and after 3 days the living fluorescent neurons were counted. The results are disclosed as a percentage of initial neurons. The comparison of survival-promoting activity of Atto-647-C-CDNF to the effect of unlabeled CDNF showed that Atto-647-labeling did not interfere with the neuroprotective activity of C-CDNF.

#### Collection of animal tissue for ELISA

Mice were sedated with terminal anaesthesia, then a cardiac puncher was performed to take a blood serum sample. Immediately after taking a blood sample, mice were transcardially perfused with cold PBS until the liver was clear from blood. The mice were then dissected on ice taking the brain first, and then the spine. These tissues were snap frozen and stored at -80 °C until use. Under dissection the brain was checked for signs of total PBS perfusion to ensure no blood remained. The left hemisphere of the brain was dissected along with the lumbar area of the spinal cord and was homogenised with or in-house lysis buffer, using a Precellys machine. The homogenate was then centrifuged, and the soluble fraction remained. The samples were diluted and analysed for C-CDNF protein concentration using an in-house ELISA.

#### Enzyme-linked immunosorbent assay (ELISA)

Recovered concentration of subcutaneously injected C-CDNF peptide from blood sera, lumbar spinal cord, and brain from WT mice were analysed using in-house-built C-CDNF ELISA. A Nunc Maxisorp immunosorbent 96-well plate was coated overnight at +4 °C with rabbit polyclonal antibody to human CDNF (Icosagen, 300-100) at 1 μg/ml in coating buffer (15mM sodium carbonate, 36mM sodium bicarbonate, pH 9.5). The following day, the plate was blocked by rinsing 3 times with ELISA Assay Buffer (Invitrogen, DS98200). For the standard curve, C-CDNF peptide was diluted in Assay Buffer in a serial dilution from 1 ng/ml to 15.6 pg/ml a blank sample was added of only Assay Buffer. All samples were diluted with Assay Buffer, applied to the plate in duplicate and incubated overnight at +4 °C with agitation. The following day, the plate was washed 3 times with Wash Buffer (WB) (0.05% Tween-20 in PBS), then detection antibody applied at 0.2 µg/ml diluted in Assay Buffer and incubated at room temperature for 2 h with agitation. After 3 washes with WB, secondary antibody polyclonal goat anti-rabbit immunoglobulins/HRP (1:1000; Agilent, P0448) diluted in Assay Buffer incubated 1.5 h at room temperature with agitation. After three washes with WB, the plate was incubated with 3,3’,5,5’-tetramethylbenzidine (TMB substrate) according to manufacturer instructions (Substrate reagent pack, DY999, R&D Systems) and absorbance was read at 450 nm minus background 540 nm (VICTOR^3^ spectrophotometer, Perkin Elmer). All reagents, antibodies and samples were applied to the plate at 100 μl. Wash buffer and Assay buffer blocking were performed with 200 µl volumes. Protein concentration of the samples was measured using DC Protein Assay (5000116, Bio-Rad) following the manufactures protocol.

### Studying the effect of C-CDNF in the 6-OHDA rat model of Parkinson’s disease

#### Stereotaxic surgeries

Stereotaxic surgeries were done under isoflurane anesthesia (4.5 % during induction, 2–3 % during maintenance). 0.1 ml of lidocaine (10 mg/ml, Orion Pharma) was injected under the scalp for local anesthesia. An incision was made to expose the skull and a high-speed drill was used to make holes in the skull. A 33G steel needle with a 10 μl syringe (Nanofil, World Precision Instruments) was used to inject 6-OHDA and C-CDNF at a 10° angle. After the injection, the needle was kept in place for 5 min to minimize backflow. Carprofen (5 mg/kg, s.c., Rimadyl, Pfizer) was used for postoperative pain relief.

Stereotaxic injections of 6-OHDA were performed as described earlier (Penttinen et al. 2016). Rats received unilateral injections of 6-OHDA hydrochloride (Sigma Aldrich) dissolved in degassed saline with 0.02 % ascorbic acid. A total of 6 µg of 6-OHDA (calculated as free base) was injected into three sites (2 µg / 1.5 µl each) at a flow rate of 0.5 µl/min in the right striatum using the following coordinates relative to the bregma: A/P +1.6, L/M +2.8, D/V −6; A/P 0.0, L/M +4.1, D/V −5.5 and A/P −2.1, L/M +4.5, D/V −5.5.

#### Administration of C-CDNF

*Intrastriatal administration*: Human C-CDNF, produced in CHO cells (Icosagen), or vehicle (PBS) was intrastriatally injected, in three locations, to 6-OHDA lesioned rats two weeks after lesioning using the same stereotaxic coordinates as with 6-OHDA injections. A total of 3.74 µg of C-CDNF was injected to the right striatum with injection volume 2 µl at a flow rate of 0.5 µl/min. *Subcutaneous administration*: C-CDNF, 50 µg or 150 µg per injection, or vehicle (PBS) was subcutaneously injected starting two weeks after 6-OHDA lesioning. Injections were given two times a week for three weeks.

#### Rotational assay

Amphetamine-induced rotational behavior was measured using automatic rotometer bowls and RotoRat software (Med Associates, Inc). Following a habituation period of 30 min, a dose of d-amphetamine sulfate (2.5 mg/kg, s.c., Division of Pharmaceutical Chemistry, University of Helsinki, Finland) dissolved in saline was given. Rotational behavior was monitored for two hours. Complete (360°) ipsilateral turns were given positive value and complete contralateral turns were given negative value. Rats rotating <50 ipsilateral turns two weeks after 6-OHDA injections were excluded from the study.

#### Tissue collection

At the end of each experiment, rats were anesthetized with an overdose of sodium pentobarbital (90 mg/kg, i.p., Orion Pharma) and perfused transcardially with PBS followed by 4 % PFA. The brains were removed, postfixed overnight in 4 % PFA, and stored in PBS containing 20 % sucrose at 4°C.

The brains were frozen and cut into 40 µm-thick coronal sections on a cryostat (Leica Instruments) in series of six. The sections were stored in a cryopreservant solution (30 % ethyleneglycol and 30 % glycerol in 0.5 M phosphate buffer) at −20°C.

#### TH DAB staining

Free-floating sections from 6-OHDA-treated rats were processed for TH-immunohistochemistry as described earlier (Albert et al. 2019). Briefly, sections were rinsed with PBS for 3x10 min, endogenous peroxidase was blocked with 0.3 % hydrogen peroxide in PBS for 30 min, sections were again rinsed with PBS and incubated in blocking buffer (4 % bovine serum albumin and 0.3 % Triton X-100 in PBS) for 1 h, then incubated overnight with the primary antibody (mouse anti-TH monoclonal, 1:2000, MAB318, Millipore) in blocking buffer at 4°C. Sections were rinsed with PBS and incubated with the secondary antibody (horse anti-mouse IgG antibody, biotinylated, 1:200, BA- 2000, Vector Laboratories) in blocking buffer for 1 h at room temperature. After rinsing with PBS, sections were incubated in avidin-biotinylated horseradish peroxidase (ABC Kit, Vector Laboratories) in PBS for 1 h, rinsed again, and finally, the staining was visualized with 3,3- diaminobenzidine (DAB) solution (DAB peroxidase substrate kit, SK-4100, Vector Laboratories) according to the manufacturer’s instructions. After rinsing with PBS, sections were placed on glass microscope slides and the slides were allowed to dry at room temperature. Lastly, sections were dehydrated with increasing ethanol series and xylene, and coverslipped with Coverquick 2000 mounting medium.

#### TH-positive cell counting

Brain sections were digitized using automated whole slide scanner (Pannoramic 250 Flash II, 3D Histech) with extended focus at a resolution of 0.22 μm/pixel. The number of TH-positive cells in the SNpc was counted using a deep convolutional neural network algorithm in Aiforia™ platform (Aiforia Technologies Oy, Helsinki, Finland) as described previously (Penttinen et al. 2018). The data are presented as percentage of the intact side that was defined as 100 %.

#### TH, Iba1, and XBP1s immunofluorescence staining

Free-floating sections were rinsed with PBS for 3x10 min, heated in 10 mM citrate buffer, pH 6, with 0.05 % Tween 20 at +80°C for 30 min, rinsed with PBS, incubated in blocking solution (4 % BSA and 0.3 % Triton X-100 in PBS) for 1 h, and transferred to the primary antibody solution (sheep anti-TH 1:2000, AB152, Sigma-Aldrich; mouse anti-Iba1 1:1000, MABN92, Sigma-Aldrich; rabbit anti-XBP-1s 1:500, E9V3E, Cell Signaling). After overnight incubation with the primary antibodies, sections were rinsed with PBS, incubated with the secondary antibodies (donkey anti-sheep IgG Alexa Fluor 488, A11015, for TH; donkey anti-mouse IgG Alexa Fluor 568, A10037, for Iba1; donkey anti-rabbit IgG Alexa Fluor 647, A31573, for XBP1s, dilution 1:1000, Invitrogen) in blocking solution for 1 h at room temperature, rinsed with PBS, transferred to DAPI (Thermo Fisher scientific, 1:2000, Cat. 62247) for 10 min at room temperature, rinsed with PBS, placed on glass slides and coverlipped with Immu-Mount (Thermo Fisher Scientific).

Laser scanning confocal micrographs of the TH+Iba1+XBP1s+ fluorescently labeled brain sections were taken with the Stellaris 8 FALCON (Leica) confocal microscope using a HC PL APO 20x/0.75 CS2 air objective. Z-stacks of 10 images at 2 μm intervals were taken from the stained area and combined into a maximum intensity projection using the imaging software Las X (version 4.4.0, Leica).

Image analysis was performed using Fiji image J (version 1.53). Iba1+ surface area was calculated from region of substantia nigra and reported as percentage of Iba1+ pixels. XBP1s+ puncta were counted manually and reported as number of positively stained aggregates localized inside Iba1+ cells.

### SOD1-G93A mouse model

Both male and female mice were used in this study. Male SOD1-G93A mice (B6SJL- Tg(SOD1*G93A)1Gur/J; Strain #:002726; RRID:IMSR_JAX:002726) were bred with female wild type mice (B6SJLF1/J Strain #:100012; RRID:IMSR_JAX:100012) both from The Jackson Laboratory within our in-house facilities. Wild type littermates were used as controls in this study. SOD1-G93A mice express the human mutant (glycine at codon 93 to alanine) superoxide dismutase 1 gene, driven by the human *SOD1* promoter. Transgenic expression was assessed by DNA ear punch test PCR (SOD1 fwd: 5’-CAC GTG GGC TCC AGC ATT-3’; SOD1 rev: 5’-TCA CCA GTC ATT TCT GCC TTT G-3’). Mice were housed according to standard conditions with mixed wild type and transgenic littermates. Mice had access to food and water ad libitum with a 12-hour dark/light cycle. Animals’ clinical status and health status were monitored daily. Recording of clinical score commenced at pre-symptomatic (11 weeks of age) until the endpoint (clinical score of 2). Body weight changes and Rotarod scoring was also done once per week on the same day before, subcutaneous injection. A clinical score scale from 5 to 2 was used to determine disease progression, according to Amyotrophic Lateral Sclerosis Therapy Development Institute (ALSTDI) guidelines. Animals were sacrificed at our predefined endpoint (clinical score of 2) when animals were no longer able to propel themselves forward with their hind limbs.

#### Analysis of motor behaviour and immunohistochemistry of SOD1-G93A mice

Motor behaviour of SOD1-G93A mice was assessed using rotarod. *<u>Accelerating rotarod</u>*: mice were tested for their ability to maintain balance on a rotating rod with accelerating speed (4-40 rpm) and the latency to fall was measured (cut off time 4 minutes) (Rota-Rod, Ugo Basile, Italy).

For immunohistochemical analyses, animals were perfused transcardially with cold PBS and then with 4% PFA; spinal cords and brains were collected and post-fixed overnight in 4% PFA.

*<u>ChAT DAB</u>.* Ten-micrometer-thick paraffin sections were cut through the spinal cord (L3-L5). Spinal cord sections were incubated with goat anti-ChAT (1:100, Merk Millipore, AB144P, MA, USA) antibody, followed by incubation with biotinylated secondary goat anti-rabbit (1:200, Vector laboratories Inc., Burlingame, USA) antibody. Antibodies were combined with standard avidin-biotin complex (ABC) reagents (Vectastain Elite ABC kit, Vector laboratories Inc., Burlingame, USA) and the signal was visualized with 3,3′-diaminobenzidine (DAB) (Sk-4100). For the quantification of ChAT^+^ neurons, the number of MNs was counted from serial lumbar spinal cord sections. The modal average of 8 sections were analysed, and the mean of counts on both sides of the spinal cord was statistically evaluated.

*<u>GFAP, IBA1 and ChAT IF.</u>* For immunohistochemistry staining of GFAP (1:2500, Neuromics- RA22101) and IBA1 (1:1000, Sigma Aldrich MABN92), 10 μm paraffin sections were deparaffinized. Heat-mediated antigen retrieval was performed using 10 mM sodium citrate buffer, pH 6. The endogenous peroxidases were inactivated followed by 1 h of blocking in TBS-T (tris-buffered saline + 0.1 % tween 20) containing 10% horse serum (HS). The sections were incubated overnight with the primary antibody at 4° C. Sections were then washed with TBS-T and the proper secondary antibody was applied for 1 hour at room temperature and coverslipped.

Laser scanning confocal micrographs of fluorescently labelled sections were taken with the Stellaris 8 FALCON (Leica, Wetzlar, Germany) confocal microscope using a HC PL APO 20x/0.75 CS2 air objective. Z-stacks were taken from ventral horn grey matter in mouse spinal cord sections. Micrographs were then combined into a maximum intensity projection using the imaging software Las X (version 4.4.0, Leica). Image analysis was performed using Fiji image J (version 1.53). Iba1+ & GFAP+ surface areas were calculated from thresholded images and reported as percentage of positive pixels. In rat brain sections, XBP1s+ & p-eIF2α+ puncta were counted manually and reported as number of positively stained aggregates localized inside Iba1+ cells.

### Statistical analysis

#### Cell and BBB penetration assays

All statistics performed in graphpad prism 9.2.0 Iodinated peptide assays: unpaired two tailed t-test Atto labelled peptide assay: Kruskal-Wallis test with Dunn’s multiple comparisons test

#### PD animal model

All statistics performed in graphpad prism 9.2.0 Amphetamine-induced rotations: Repeated measures ANOVA with Dunnett’s multiple comparisons test. Area under curve followed by unpaired two tailed t-test or one-way ANOVA with Tukey’s multiple comparisons test.

Immunofluorescence and immunohistochemistry: unpaired two tailed t-test or one-way ANOVA followed by Tukey’s multiple comparisons test.

Grubb’s test was used to identify outliers.

#### ALS animal model

All statistics performed in graphpad prism 9.2.0

Clinical score: Mann-Whitney test with Holm-Šídák method Rotarod: Mann-Whitney tests with Holm-Šídák method

Immunofluorescence and immunohistochemistry: unpaired two tailed t-test

## Supplementary methods

### Primary cultures of sympathetic neurons and microinjection

Culture of mouse superior cervical ganglion (SCG) sympathetic neurons and microinjection of these neurons was performed as described earlier (Yu et al. 2003). Briefly, the neurons of postnatal day 1– 2 NMRI strain mice were grown 6 DIV on polyornithine-laminin (P3655 and CC095, Sigma-Aldrich)–coated dishes or glass coverslips with 30 ng/ml of 2.5 S mouse NGF (G5141, Promega) in the Neurobasal medium containing B27 supplement (17504044, Invitrogen). The nuclei were then microinjected with the expression plasmid for CDNF or C-CDNF together with a reporter plasmid for enhanced green fluorescent protein (EGFP), at concentration of 10 ng/μl in each experiment. For protein microinjection, recombinant CDNF, C-CDNF protein or C-CDNF peptide in PBS at 200ng/ul was microinjected directly into the cytoplasm together with fluorescent reporter Dextran Texas Red (MW 70000 Da) (D1864, Invitrogen, Molecular Probes) that facilitates identification of successfully injected neurons. Next day, tunicamycin (2 µM) (ab120296, Abcam) was added and living fluorescent (EGFP-expressing or Dextran Texas Red-containing) neurons were counted three days later and expressed as percentage of initial living fluorescent neurons counted 2–3 hours after microinjection.

---

**Supplement figure 1.**
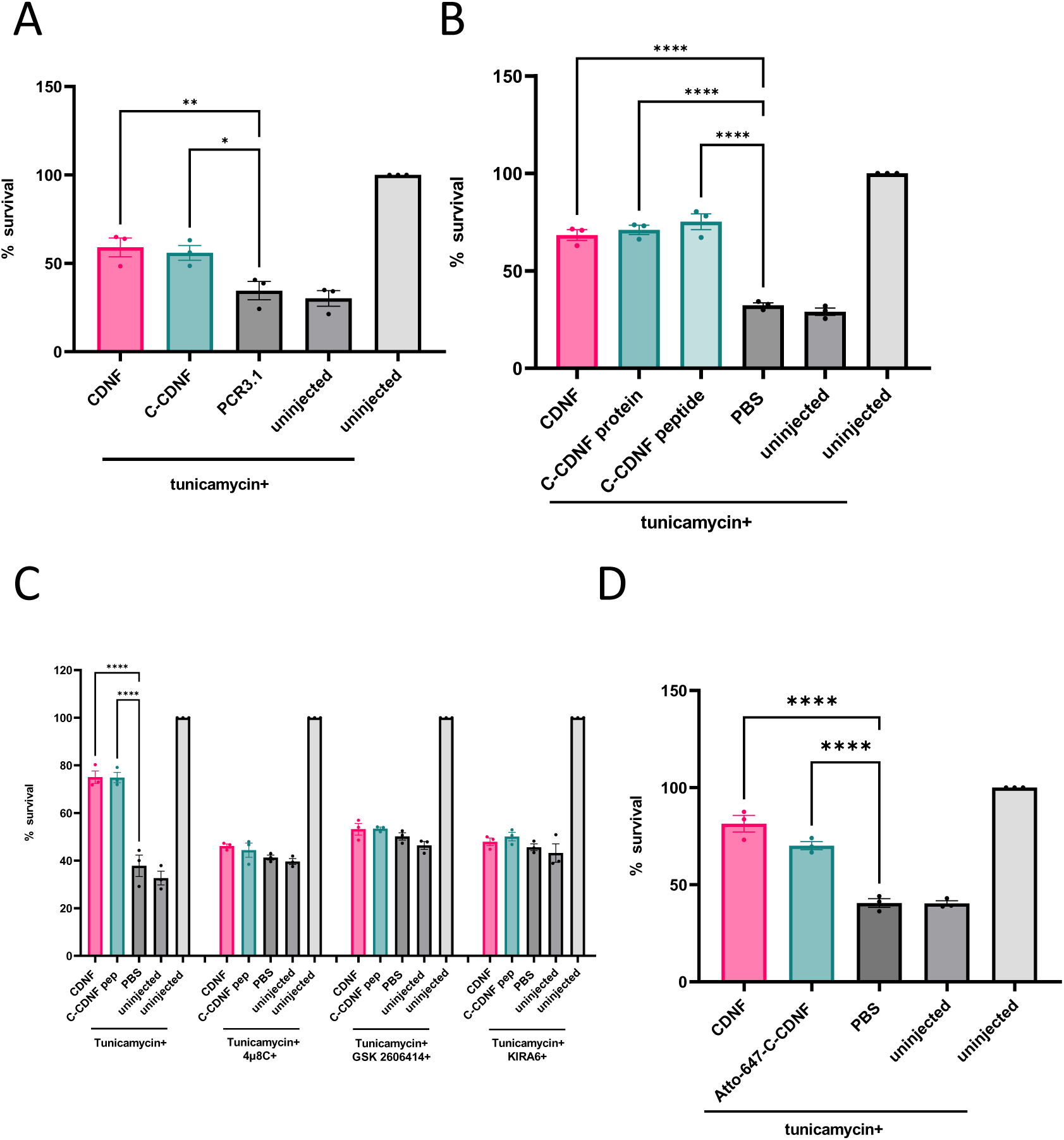
C-CDNF promotes survival of ER–stressed SCG sympathetic neurons by regulating UPR signaling. (A-B) Microinjection with C-CDNF expression plasmid (A) or recombinant C-CDNF protein or chemically synthesized peptide (B) protected mouse SCG neurons from tunicamycin (2µM) treatment. After 72 hours, the number of living and fluorescent neurons was counted and expressed as % of initially injected neurons. Shown are means of 3 experiments ± S.D. (C) Survival-promoting activity of C-CDNF against ER stress is dependent on the activation of IRE1α and PERK pathways. Mouse SCG neurons were microinjected with C-CDNF peptide and then treated with tunicamycin together with PERK inhibitor GSK2606414 or IREα inhibitors 4μ8C or KIRA6. Shown are the means of 3 experiments ± S.D. (D) Atto-647-C-CDNF is biologically active. Survival percentages of mouse SCG neurons in C-CDNF plasmid- or protein-injected groups were compared to the empty vector or PBS-injected controls of the same treatment group using ordinary one-way analysis of variance (ANOVA) and Sidak’s multiple comparison post hoc test. *, **, ***, ****p < 0.05, p < 0.01, p < 0.001, p < 0.0001 in A-C.

**Table S1.**
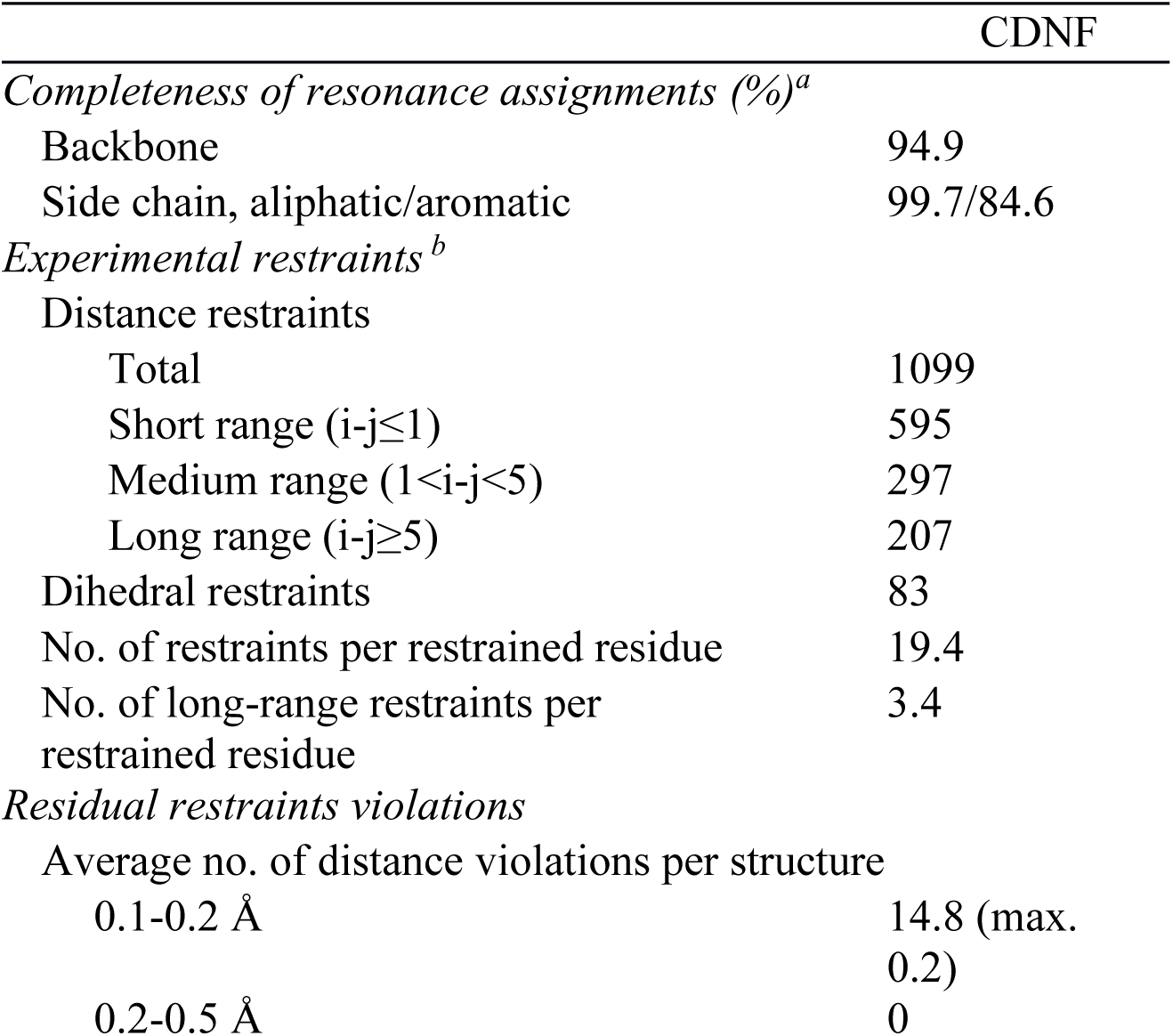
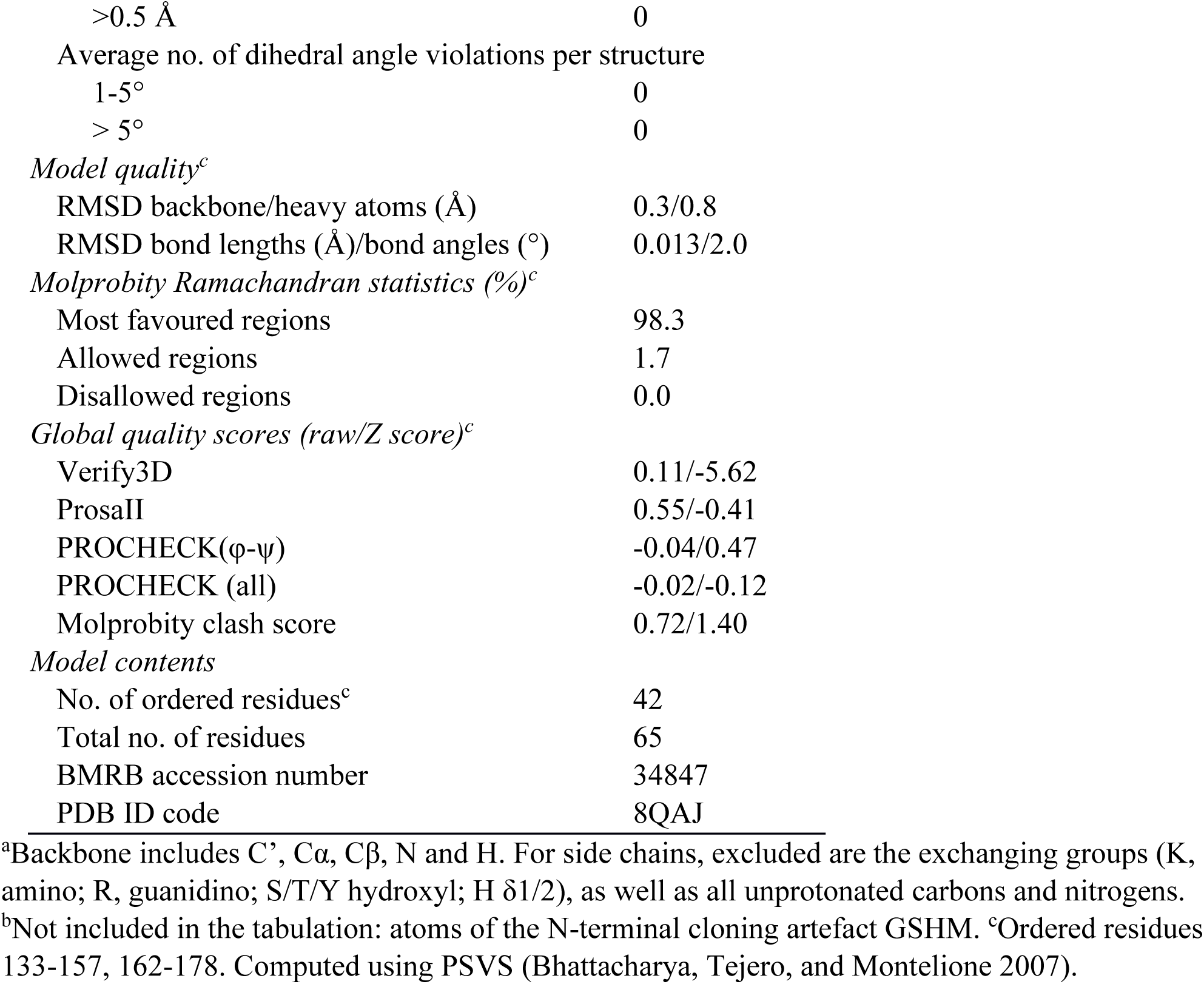
NMR restraints and structural statistics for CDNF. Ensemble contains 20 conformers of least restraint violations.

## REFERENCES

1. Airaksinen MS, Saarma M. The GDNF family: signalling, biological functions and therapeutic value. Nat Rev Neurosci. 2002 May;3(5):383–94. doi: 10.1038/nrn812. PMID: 11988777.

2. Airavaara M, Harvey BK, Voutilainen MH, Shen H, Chou J, Lindholm P, Lindahl M, Tuominen RK, Saarma M, Hoffer B, Wang Y. CDNF protects the nigrostriatal dopamine system and promotes recovery after MPTP treatment in mice. Cell Transplant. 2012;21(6):1213–23. doi: 10.3727/096368911X600948. Epub 2011 Sep 22. PMID: 21943517; PMCID: PMC3753365.

3. Albert, K., D. P. Raymundo, A. Panhelainen, A. Eesmaa, L. Shvachiy, G. R. Araujo, P. Chmielarz, X. Yan, A. Singh, Y. Cordeiro, F. L. Palhano, D. Foguel, K. C. Luk, A. Domanskyi, M. H. Voutilainen, H. J. Huttunen, T. F. Outeiro, M. Saarma, M. S. Almeida, and M. Airavaara. 2021. ’Cerebral dopamine neurotrophic factor reduces alpha-synuclein aggregation and propagation and alleviates behavioral alterations in vivo’, Mol Ther, 29: 2821–40.

4. Albert, K., M. H. Voutilainen, A. Domanskyi, T. P. Piepponen, S. Ahola, R. K. Tuominen, C. Richie, B. K. Harvey, and M. Airavaara. 2019. ’Downregulation of tyrosine hydroxylase phenotype after AAV injection above substantia nigra: Caution in experimental models of Parkinson’s disease’, J Neurosci Res, 97: 346–61.

5. Apostolou A, Shen Y, Liang Y, Luo J, Fang S. Armet, a UPR-upregulated protein, inhibits cell proliferation and ER stress-induced cell death. Exp Cell Res. 2008,314(13):2454-67. doi: 10.1016/j.yexcr.2008.05.001. Epub 2008 May 13. PMID: 18561914; PMCID: PMC6719340.

6. Arancibia D, Zamorano P, Andrés ME. CDNF induces the adaptive unfolded protein response and attenuates endoplasmic reticulum stress-induced cell death. Biochim Biophys Acta Mol Cell Res. 2018, 1865(11 Pt A):1579-1589. doi: 10.1016/j.bbamcr.2018.08.012. Epub 2018 Aug 23. PMID: 30327199.

7. Axten, J. M., J. R. Medina, Y. Feng, A. Shu, S. P. Romeril, S. W. Grant, W. H. Li, D. A. Heerding, E. Minthorn, T. Mencken, C. Atkins, Q. Liu, S. Rabindran, R. Kumar, X. Hong, A. Goetz, T. Stanley, J. D. Taylor, S. D. Sigethy, G. H. Tomberlin, A. M. Hassell, K. M. Kahler, L. M. Shewchuk, and R. T. Gampe. 2012. ’Discovery of 7-methyl-5-(1-[3- (trifluoromethyl)phenyl]acetyl-2,3-dihydro-1H-indol-5-yl)-7H-pyrrolo[2,3-d]pyrimidin-4-amine (GSK2606414), a potent and selective first-in-class inhibitor of protein kinase R (PKR)-like endoplasmic reticulum kinase (PERK)’, J Med Chem, 55: 7193–207.

8. Barker, R. A., A. Bjorklund, D. M. Gash, A. Whone, A. Van Laar, J. H. Kordower, K. Bankiewicz, K. Kieburtz, M. Saarma, S. Booms, H. J. Huttunen, A. P. Kells, M. S. Fiandaca, A. J. Stoessl, D. Eidelberg, H. Federoff, M. H. Voutilainen, D. T. Dexter, J. Eberling, P. Brundin, L. Isaacs, L. Mursaleen, E. Bresolin, C. Carroll, A. Coles, B. Fiske, H. Matthews, C. Lungu, R. K. Wyse, S. Stott, and A. E. Lang. 2020. ’GDNF and Parkinson’s Disease: Where Next? A Summary from a Recent Workshop’, J Parkinsons Dis, 10: 875–91.

9. Bhattacharya, A., R. Tejero, and G. T. Montelione. 2007. ’Evaluating protein structures determined by structural genomics consortia’, Proteins, 66: 778–95.

10. Bäck S, Peränen J, Galli E, Pulkkila P, Lonka-Nevalaita L, Tamminen T, Voutilainen MH, Raasmaja A, Saarma M, Männistö PT, Tuominen RK. Gene therapy with AAV2-CDNF provides functional benefits in a rat model of Parkinson’s disease. Brain Behav. 2013 Mar;3(2):75–88. doi: 10.1002/brb3.117. Epub 2013 Jan 14. PMID: 23532969; PMCID: PMC3607149

11. Bradley LH, Fuqua J, Richardson A, Turchan-Cholewo J, Ai Y, Kelps KA, Glass JD, He X, Zhang Z, Grondin R, Littrell OM, Huettl P, Pomerleau F, Gash DM, Gerhardt GA. Dopamine neuron stimulating actions of a GDNF propeptide. PLoS One. 2010 Mar 18;5(3):e9752. doi: 10.1371/journal.pone.0009752. PMID: 20305789; PMCID: PMC2841203.

12. Cao, D., X. Ma, J. Cai, J. Luan, A. J. Liu, R. Yang, Y. Cao, X. Zhu, H. Zhang, Y. X. Chen, Y. Shi, G. X. Shi, D. Zou, X. Cao, M. J. Grusby, Z. Xie, and W. J. Zhang. 2016. ’ZBTB20 is required for anterior pituitary development and lactotrope specification’, Nat Commun, 7: 11121.

13. Cross, B. C., P. J. Bond, P. G. Sadowski, B. K. Jha, J. Zak, J. M. Goodman, R. H. Silverman, T. A. Neubert, I. R. Baxendale, D. Ron, and H. P. Harding. 2012. ’The molecular basis for selective inhibition of unconventional mRNA splicing by an IRE1-binding small molecule’, Proc Natl Acad Sci U S A, 109: E869–78.

14. Danilova, T., I. Belevich, H. Li, E. Palm, E. Jokitalo, T. Otonkoski, and M. Lindahl. 2019. ’MANF Is Required for the Postnatal Expansion and Maintenance of Pancreatic beta-Cell Mass in Mice’, Diabetes, 68: 66–80.

15. De Lorenzo, F., P. Luningschror, J. Nam, L. Beckett, F. Pilotto, E. Galli, P. Lindholm, C. Rudt von Collenberg, S. T. Mungwa, S. Jablonka, J. Kauder, N. Thau-Habermann, S. Petri, D. Lindholm, S. Saxena, M. Sendtner, M. Saarma, and M. H. Voutilainen. 2023. ’CDNF rescues motor neurons in models of amyotrophic lateral sclerosis by targeting endoplasmic reticulum stress’, Brain, 146: 3783–99.

16. Delaglio, F., S. Grzesiek, G. W. Vuister, G. Zhu, J. Pfeifer, and A. Bax. 1995. ’NMRPipe: a multidimensional spectral processing system based on UNIX pipes’, J Biomol NMR, 6: 277–93.

17. Eesmaa, A., L. Y. Yu, H. Goos, T. Danilova, K. Noges, E. Pakarinen, M. Varjosalo, M. Lindahl, P. Lindholm, and M. Saarma. 2022. ’CDNF Interacts with ER Chaperones and Requires UPR Sensors to Promote Neuronal Survival’, Int J Mol Sci, 23.

18. Eesmaa, A., L. Y. Yu, H. Goos, K. Noges, V. Kovaleva, M. Hellman, R. Zimmermann, M. Jung, P. Permi, M. Varjosalo, P. Lindholm, and M. Saarma. 2021. ’The cytoprotective protein MANF promotes neuronal survival independently from its role as a GRP78 cofactor’, J Biol Chem, 296: 100295.

19. Galli, E., A. Planken, L. Kadastik-Eerme, M. Saarma, P. Taba, and P. Lindholm. 2019. ’Increased Serum Levels of Mesencephalic Astrocyte-Derived Neurotrophic Factor in Subjects With Parkinson’s Disease’, Front Neurosci, 13: 929.

20. Ghosh, R., L. Wang, E. S. Wang, B. G. Perera, A. Igbaria, S. Morita, K. Prado, M. Thamsen, D. Caswell, H. Macias, K. F. Weiberth, M. J. Gliedt, M. V. Alavi, S. B. Hari, A. K. Mitra, B. Bhhatarai, S. C. Schurer, E. L. Snapp, D. B. Gould, M. S. German, B. J. Backes, D. J. Maly, S. A. Oakes, and F. R. Papa. 2014. ’Allosteric inhibition of the IRE1alpha RNase preserves cell viability and function during endoplasmic reticulum stress’, Cell, 158: 534–48.

21. Guntert, P., and L. Buchner. 2015. ’Combined automated NOE assignment and structure calculation with CYANA’, J Biomol NMR, 62: 453–71.

22. Hellman, M., U. Arumae, L. Y. Yu, P. Lindholm, J. Peranen, M. Saarma, and P. Permi. 2011. ’Mesencephalic astrocyte-derived neurotrophic factor (MANF) has a unique mechanism to rescue apoptotic neurons’, J Biol Chem, 286: 2675–80.

23. Hetz, C., and B. Mollereau. 2014. ’Disturbance of endoplasmic reticulum proteostasis in neurodegenerative diseases’, Nat Rev Neurosci, 15: 233–49.

24. Hetz, C., and S. Saxena. 2017. ’ER stress and the unfolded protein response in neurodegeneration’, Nat Rev Neurol, 13: 477–91.

25. Huttunen, H. J., S. Booms, M. Sjogren, V. Kerstens, J. Johansson, R. Holmnas, J. Koskinen, N. Kulesskaya, P. Fazio, M. Woolley, A. Brady, J. Williams, D. Johnson, N. Dailami, W. Gray, R. Levo, M. Saarma, C. Halldin, J. Marjamaa, J. Resendiz-Nieves, I. Grubor, G. Lind, J. Eerola-Rautio, T. Mertsalmi, M. Andreasson, G. Paul, J. Rinne, R. Kivisaari, H. Bjartmarz, P. Almqvist, A. Varrone, F. Scheperjans, H. Widner, and P. Svenningsson. 2023. ’Intraputamenal Cerebral Dopamine Neurotrophic Factor in Parkinson’s Disease: A Randomized, Double-Blind, Multicenter Phase 1 Trial’, Mov Disord, 38: 1209–22.

26. Kikuchi, H., G. Almer, S. Yamashita, C. Guegan, M. Nagai, Z. Xu, A. A. Sosunov, G. M. McKhann, 2nd, and S. Przedborski. 2006. ’Spinal cord endoplasmic reticulum stress associated with a microsomal accumulation of mutant superoxide dismutase-1 in an ALS model’, Proc Natl Acad Sci U S A, 103: 6025-30.

27. Kovaleva, V., L. Y. Yu, L. Ivanova, O. Shpironok, J. Nam, A. Eesmaa, E. P. Kumpula, S. Sakson, U. Toots, M. Ustav, J. T. Huiskonen, M. H. Voutilainen, P. Lindholm, M. Karelson, and M. Saarma. 2023. ’MANF regulates neuronal survival and UPR through its ER-located receptor IRE1alpha’, Cell Rep, 42: 112066.

28. Latge, C., K. M. Cabral, G. A. de Oliveira, D. P. Raymundo, J. A. Freitas, L. Johanson, L. F. Romao, F. L. Palhano, T. Herrmann, M. S. Almeida, and D. Foguel. 2015. ’The Solution Structure and Dynamics of Full-length Human Cerebral Dopamine Neurotrophic Factor and Its Neuroprotective Role against alpha-Synuclein Oligomers’, J Biol Chem, 290: 20527–40.

29. Lindahl, M., T. Danilova, E. Palm, P. Lindholm, V. Voikar, E. Hakonen, J. Ustinov, J. O. Andressoo, B. K. Harvey, T. Otonkoski, J. Rossi, and M. Saarma. 2014. ’MANF is indispensable for the proliferation and survival of pancreatic beta cells’, Cell Rep, 7: 366–75.

30. Lindholm, P., and M. Saarma. 2022. ’Cerebral dopamine neurotrophic factor protects and repairs dopamine neurons by novel mechanism’, Mol Psychiatry, 27: 1310–21.

31. Lindholm, P., M. H. Voutilainen, J. Lauren, J. Peranen, V. M. Leppanen, J. O. Andressoo, M. Lindahl, S. Janhunen, N. Kalkkinen, T. Timmusk, R. K. Tuominen, and M. Saarma. 2007. ’Novel neurotrophic factor CDNF protects and rescues midbrain dopamine neurons in vivo’, Nature, 448: 73–7.

32. Liu, H., C. Zhao, L. Zhong, J. Liu, S. Zhang, B. Cheng, and L. Gong. 2015. ’Key subdomains in the C-terminal of cerebral dopamine neurotrophic factor regulate the protein secretion’, Biochem Biophys Res Commun, 465: 427–32.

33. Mahato, A. K., J. Kopra, J. M. Renko, T. Visnapuu, I. Korhonen, N. Pulkkinen, M. M. Bespalov, A. Domanskyi, E. Ronken, T. P. Piepponen, M. H. Voutilainen, R. K. Tuominen, M. Karelson, Y. A. Sidorova, and M. Saarma. 2020. ’Glial cell line-derived neurotrophic factor receptor Rearranged during transfection agonist supports dopamine neurons in Vitro and enhances dopamine release In Vivo’, Mov Disord, 35: 245–55.

34. Manfredsson FP, Polinski NK, Subramanian T, Boulis N, Wakeman DR, Mandel RJ. The Future of GDNF in Parkinson’s Disease. Front Aging Neurosci. 2020 Dec 7;12:593572. doi: 10.3389/fnagi.2020.593572. PMID: 33364933; PMCID: PMC7750181.

35. Matlik, K., L. Y. Yu, A. Eesmaa, M. Hellman, P. Lindholm, J. Peranen, E. Galli, J. Anttila, M. Saarma, P. Permi, M. Airavaara, and U. Arumae. 2015. ’Role of two sequence motifs of mesencephalic astrocyte-derived neurotrophic factor in its survival-promoting activity’, Cell Death Dis, 6: e2032.

36. McQuin, C., A. Goodman, V. Chernyshev, L. Kamentsky, B. A. Cimini, K. W. Karhohs, M. Doan, L. Ding, S. M. Rafelski, D. Thirstrup, W. Wiegraebe, S. Singh, T. Becker, J. C. Caicedo, and A. E. Carpenter. 2018. ’CellProfiler 3.0: Next-generation image processing for biology’, PLoS Biol, 16: e2005970.

37. Nadella, R., M. H. Voutilainen, M. Saarma, J. A. Gonzalez-Barrios, B. A. Leon-Chavez, J. M. Jimenez, S. H. Jimenez, L. Escobedo, and D. Martinez-Fong. 2014. ’Transient transfection of human CDNF gene reduces the 6-hydroxydopamine-induced neuroinflammation in the rat substantia nigra’, J Neuroinflammation, 11: 209.

38. Nagai, M., D. B. Re, T. Nagata, A. Chalazonitis, T. M. Jessell, H. Wichterle, and S. Przedborski. 2007. ’Astrocytes expressing ALS-linked mutated SOD1 release factors selectively toxic to motor neurons’, Nat Neurosci, 10: 615–22.

39. Norisada, J., Y. Hirata, F. Amaya, K. Kiuchi, and K. Oh-hashi. 2016. ’A Comparative Analysis of the Molecular Features of MANF and CDNF’, PLoS One, 11: e0146923.

40. Parkash, V., P. Lindholm, J. Peranen, N. Kalkkinen, E. Oksanen, M. Saarma, V. M. Leppanen, and A. Goldman. 2009. ’The structure of the conserved neurotrophic factors MANF and CDNF explains why they are bifunctional’, Protein Eng Des Sel, 22: 233–41.

41. Penttinen, A. M., I. Suleymanova, K. Albert, J. Anttila, M. H. Voutilainen, and M. Airavaara. 2016. ’Characterization of a new low-dose 6-hydroxydopamine model of Parkinson’s disease in rat’, J Neurosci Res, 94: 318–28.

42. Permi, P., and A. Annila. 2001. ’A new approach for obtaining sequential assignment of large proteins’, J Biomol NMR, 20: 127–33.

43. Permi P, Annila A. 2004. Coherence transfer in proteins. Prog Nucl Magn Reson Spectrosc 44:97– 13

44. Permi, P., H. Tossavainen, and M. Hellman. 2004. ’Efficient assignment of methyl resonances: enhanced sensitivity by gradient selection in a DE-MQ-(H)CC(m)Ht (m)-TOCSY experiment’, J Biomol NMR, 30: 275–82.

45. Petrova, P., A. Raibekas, J. Pevsner, N. Vigo, M. Anafi, M. K. Moore, A. E. Peaire, V. Shridhar, D. I. Smith, J. Kelly, Y. Durocher, and J. W. Commissiong. 2003. ’MANF: a new mesencephalic, astrocyte-derived neurotrophic factor with selectivity for dopaminergic neurons’, J Mol Neurosci, 20: 173–88.

46. Saarma M, Airavaara, M., Voutilainen, M.H., Yu, L.Y., and Lindahl, M. (2018) C-terminal cdnf and manf fragments, pharmaceutical compositions comprising same and uses thereof. Patent. Publication number: 20200071372

47. Sattler M, Schleucher J, Griesinger C (1999) Heteronuclear multidimensional NMR experiments for the structure determination of proteins in solution employing pulsed field gradients. Prog Nucl Magn Reson Spectrosc 34:93–202.

48. Sehnal, D, Bittrich, S, Deshpande, M, Svobodová, R, Berka, K, Bazgier, V, et al. Mol* viewer: modern web app for 3D visualization and analysis of large biomolecular structures. Nucleic Acids Res. (2021) 49:W431–7. doi: 10.1093/nar/gkab314

49. Selvaraj BT, Livesey MR, Zhao C, Gregory JM, James OT, Cleary EM, Chouhan AK, Gane AB, Perkins EM, Dando O, Lillico SG, Lee YB, Nishimura AL, Poreci U, Thankamony S, Pray M, Vasistha NA, Magnani D, Borooah S, Burr K, Story D, McCampbell A, Shaw CE, Kind PC, Aitman TJ, Whitelaw CBA, Wilmut I, Smith C, Miles GB, Hardingham GE, Wyllie DJA, Chandran S. C9ORF72 repeat expansion causes vulnerability of motor neurons to Ca^2+^-permeable AMPA receptor-mediated excitotoxicity. Nat Commun. 2018 Jan 24;9(1):347. doi: 10.1038/s41467-017-02729-0. PMID: 29367641; PMCID: PMC5783946.

50. Shen, Y., and A. Bax. 2013. ’Protein backbone and sidechain torsion angles predicted from NMR chemical shifts using artificial neural networks’, J Biomol NMR, 56: 227–41.

51. Sidorova YA, Saarma M. Can Growth Factors Cure Parkinson’s Disease? Trends Pharmacol Sci. 2020 Dec;41(12):909–922. doi: 10.1016/j.tips.2020.09.010. Epub 2020 Oct 22. PMID: 33198924.

52. Szegezdi, E., S. E. Logue, A. M. Gorman, and A. Samali. 2006. ’Mediators of endoplasmic reticulum stress-induced apoptosis’, EMBO Rep, 7: 880–5.

53. Thoenen, H., and M. Sendtner. 2002. ’Neurotrophins: from enthusiastic expectations through sobering experiences to rational therapeutic approaches’, Nat Neurosci, 5 Suppl: 1046-50.

54. Tossavainen, H., and P. Permi. 2004. ’Optimized pathway selection in intraresidual triple-resonance experiments’, J Magn Reson, 170: 244–51.

55. Trokovic R, Weltner J, Noisa P, Raivio T, Otonkoski T. Combined negative effect of donor age and time in culture on the reprogramming efficiency into induced pluripotent stem cells. Stem Cell Res. 2015 Jul;15(1):254–62. doi: 10.1016/j.scr.2015.06.001. Epub 2015 Jun 11. PMID: 26096152.

56. Tseng, K. Y., V. Stratoulias, W. F. Hu, J. S. Wu, V. Wang, Y. H. Chen, A. Seelbach, H. J. Huttunen, N. Kulesskaya, C. Y. Pang, J. L. Chou, M. Lindahl, M. Saarma, L. C. Huang, M. Airavaara, and H. K. Liew. 2023. ’Augmenting hematoma-scavenging capacity of innate immune cells by CDNF reduces brain injury and promotes functional recovery after intracerebral hemorrhage’, Cell Death Dis, 14: 128.

57. Voutilainen, M. H., U. Arumae, M. Airavaara, and M. Saarma. 2015. ’Therapeutic potential of the endoplasmic reticulum located and secreted CDNF/MANF family of neurotrophic factors in Parkinson’s disease’, FEBS Lett, 589: 3739–48.

58. Voutilainen, M. H., S. Back, J. Peranen, P. Lindholm, A. Raasmaja, P. T. Mannisto, M. Saarma, and R. K. Tuominen. 2011. ’Chronic infusion of CDNF prevents 6-OHDA-induced deficits in a rat model of Parkinson’s disease’, Exp Neurol, 228: 99–108.

59. Voutilainen, M. H., S. Back, E. Porsti, L. Toppinen, L. Lindgren, P. Lindholm, J. Peranen, M. Saarma, and R. K. Tuominen. 2009. ’Mesencephalic astrocyte-derived neurotrophic factor is neurorestorative in rat model of Parkinson’s disease’, J Neurosci, 29: 9651–9.

60. Voutilainen, M. H., F. De Lorenzo, P. Stepanova, S. Back, L. Y. Yu, P. Lindholm, E. Porsti, M. Saarma, P. T. Mannisto, and R. K. Tuominen. 2017. ’Evidence for an Additive Neurorestorative Effect of Simultaneously Administered CDNF and GDNF in Hemiparkinsonian Rats: Implications for Different Mechanism of Action’, eNeuro, 4.

61. Vranken, W. F., W. Boucher, T. J. Stevens, R. H. Fogh, A. Pajon, M. Llinas, E. L. Ulrich, J. L. Markley, J. Ionides, and E. D. Laue. 2005. ’The CCPN data model for NMR spectroscopy: development of a software pipeline’, Proteins, 59: 687–96.

62. Barker RA, Björklund A, Gash DM, Whone A, Van Laar A, Kordower JH, Bankiewicz K, Kieburtz K, Saarma M, Booms S, Huttunen HJ, Kells AP, Fiandaca MS, Stoessl AJ, Eidelberg D, Federoff H, Voutilainen MH, Dexter DT, Eberling J, Brundin P, Isaacs L, Mursaleen L, Bresolin E, Carroll C, Coles A, Fiske B, Matthews H, Lungu C, Wyse RK, Stott S, Lang AE. GDNF and Parkinson’s Disease: Where Next? A Summary from a Recent Workshop. J Parkinsons Dis. 2020;10(3):875–891. doi: 10.3233/JPD-202004. PMID: 32508331; PMCID: PMC7458523.

63. Yu, L. Y., E. Jokitalo, Y. F. Sun, P. Mehlen, D. Lindholm, M. Saarma, and U. Arumae. 2003. ’GDNF-deprived sympathetic neurons die via a novel nonmitochondrial pathway’, J Cell Biol, 163: 987–97.

64. Yu, L. Y., M. Saarma, and U. Arumae. 2008. ’Death receptors and caspases but not mitochondria are activated in the GDNF- or BDNF-deprived dopaminergic neurons’, J Neurosci, 28: 7467–75.

65. Zhao, H., L. Cheng, X. Du, Y. Hou, Y. Liu, Z. Cui, and L. Nie. 2016. ’Transplantation of Cerebral Dopamine Neurotrophic Factor Transducted BMSCs in Contusion Spinal Cord Injury of Rats: Promotion of Nerve Regeneration by Alleviating Neuroinflammation’, Mol Neurobiol, 53: 187–99.

66. Zhao, H., L. Cheng, Y. Liu, W. Zhang, S. Maharjan, Z. Cui, X. Wang, D. Tang, and L. Nie. 2014. ’Mechanisms of anti-inflammatory property of conserved dopamine neurotrophic factor: inhibition of JNK signaling in lipopolysaccharide-induced microglia’, J Mol Neurosci, 52: 186–92.

